# DriverOmicsNet: An Integrated Graph Convolutional Network for Multi-Omics Exploration of Cancer Driver Genes

**DOI:** 10.1101/2024.07.21.604474

**Authors:** Yang-Hong Dai, Chia-Jun Chang, Po-Chien Shen, Wun-Long Jheng, Yu-Guang Chen

## Abstract

**Background:** Cancer is a complex and heterogeneous group of diseases driven by genetic mutations and molecular changes. Identifying and characterizing cancer driver genes (CDgs) is crucial for understanding cancer biology and guiding precision oncology. Integrating multi-omics data can reveal the intricate molecular interactions underlying cancer progression and treatment responses.

**Methods:** We developed a graph convolutional network (GCN) framework, DriverOmicsNet, that integrates multi-omics data using STRING protein-protein interaction (PPI) networks and correlation-based weighted correlation network analysis (WGCNA). We applied this framework to 15 cancer types, analyzing 5555 tumor samples to predict cancer-related features such as homologous recombination deficiency (HRD), cancer stemness, immune clusters, tumor stage, and survival outcomes.

**Findings:** DriverOmicsNet demonstrated superior predictive accuracy and model performance metrics across all target labels when compared with GCN models based on STRING network alone. Gene expression emerged as the most significant feature, reflecting the dynamic and functional state of cancer cells. The combined use of STRING PPI and WGCNA networks enhanced the identification of key driver genes and their interactions.

**Interpretation:** Our study highlights the effectiveness of using GCNs to integrate multi-omics data for precision oncology. The integration of STRING PPI and WGCNA networks provides a comprehensive framework that improves predictive power and facilitates the understanding of cancer biology, paving the way for more tailored treatments.

## Introduction

Cancer is a complex and heterogeneous group of diseases and continues to pose a significant challenge to medical science. The intricate web of molecular alterations within cancer cells, driven by a myriad of genetic mutations and molecular changes, has made it crucial to decipher the underlying mechanisms orchestrating the disease’s initiation, progression, and response to treatment.^1^ Among these molecular alterations, the identification and characterization of cancer driver genes (CDgs) have emerged as pivotal milestones in our quest to understand and combat cancer.^2^

Driver genes, unlike their passenger counterparts, play a crucial role in cancer development by endowing tumor cells with a selective growth advantage.^3^ Mutations in these genes initiate a cascade of molecular events, promoting uncontrolled cell growth, replicative immortality, invasion, metastasis, deregulation of energy metabolism, and evasion of immune suppression.^4^ Consequently, discerning the molecular signatures and functional implications of CDgs has become imperative in guiding precision oncology, where treatments are tailored to the unique genetic makeup of each patient’s tumor. In recent years, the advent of next-generation sequencing and high-throughput multi-omics technologies has brought us closer to comprehending the intricate landscape of cancer biology.^5^ These technologies have enabled researchers to simultaneously explore DNA mutations, gene expression, copy number variations (CNV), epigenetic modifications, protein levels, and metabolic profiles, collectively referred to as “multi-omics” data. These multi-omics data sources offer unprecedented insights into the molecular underpinnings of cancer, making it possible to uncover the complex interplay between genetic alterations, gene expression patterns, and clinical outcomes. In addition to mutations that characterize ’dysfunctional’ events in genes, other multi-omics data contribute to a holistic understanding by shedding light on ‘dysregulation’ events in oncogenesis.^6^ However, the challenge lies in integrating these diverse layers of information to elucidate the causal relationships between molecular signatures and cancer phenotypes.

Numerous approaches have emerged over the years to tackle the integration of multi-omics data. Many of these efforts have centered on unsupervised data integration, often lacking the context of sample labels.^7–9^ However, with the rise of personalized medicine, the availability of meticulously curated datasets with comprehensive sample annotations, characterizing phenotypes or traits, has expanded significantly. Consequently, there is a growing interest in supervised multi-omics integration techniques, capable of discerning disease-related biomarkers and making predictions for new samples.

In recent years, deep learning (DL) has showcased its formidable prowess in bioinformatics^10^. For example, Wang et al. applied DL algorithms, specifically leveraging graph neural networks (GNNs) to circumvent potential biases inherent in traditional methods like feature concatenation and ensemble techniques.^11^ In their pioneering work (MOGONET), they harnessed the capabilities of GNNs to uncover correlations among diverse omics data types. They adopted a novel approach by training distinct GNNs, each tailored to a specific omics data type, based on weighted similarity networks. They significantly improved patient classification, revealing pivotal biomarkers for Alzheimer’s disease and breast cancer. However, their methodology fell short in the integration of all omics data into a unified graph and only offered a datatype-specific patient embedding.^12^ To better understand how multi-omics data can elucidate relationships among CDgs, it is necessary to integrate the multi-omics features simultaneously for a local graph network. Additionally, MOGONET does not train on curated network structure such as protein-protein interaction (PPI) network, which plays essential roles in structuring and mediating biological processes.^13^

To address the aforementioned limitations, we developed a framework of graph convolutional network (GCN) named DriverOmicsNet that combined multi-omics graph embedding derived from STRING PPI network and correlation-based weighted correlation network analysis (WGCNA) for CDgs across multiple cancer types. DriverOmicsNet demonstrates remarkable predictive accuracy and excels in various model performance metrics, including tasks related to homologous DNA repair deficiency, cancer stemness, immune cluster, tumor stage, and survival prediction. Moreover, we have delved into the interpretability of our GCN models, revealing their ability to discern essential driver markers. By leveraging GNNs and a wealth of multi-omics data, our study embarks on a transformative journey that redefines the precision oncology paradigm. Our goal is to bridge the gap between the detailed molecular phenotypes of cancer cells and clinical outcomes, paving the way for tailored treatments that improve patient care.

## Methods

### Tumor samples for model construction

The multi-omics data utilized in this study were sourced from UCSC Xena platform (https://xena.ucsc.edu), a comprehensive repository of cancer genomics data. Access to the omics data was facilitated through the *UCSCXenaTools* package in R.^14^ This package enabled seamless retrieval and preprocessing of the diverse molecular datasets, ensuring the high quality and reliability of the data utilized in our study. To ensure the robustness and statistical significance of our analyses, cancer cohorts with a patient sample size of fewer than 100 individuals were excluded. Finally, an overall of 15 cancer types and 5555 tumor samples were evaluated.

### Multi-omics data type and preprocessing

### mRNA gene expression

RSEM expected count for each gene provided by the UCSC Toil Recompute Compendium were downloaded, normalized and processed according to the protocol described by Chen et al.^15,16^ Briefly, the expected counts were back transformed, followed by normalization using voom method from limma package for linear modeling.^17^ Additional 2656 cancer-free healthy tissue samples for corresponding cancer type (Genotype-Tissue Ex, GTEx) were used for control. The voom-normalized gene expression was then used for downstream WGCNA and multi-omics integration for GNN models. To further ensure the reliability of the CDgs used in our study, we employed a gene significance (GS) metric derived from WGCNA. GS was defined as the mediated p-value of each gene in the linear regression models correlating mRNA expression with the traits of interest. This method allowed us to evaluate the significance of CDgs and their associations with tumor phenotype.

### CNV and Mutation

The retrieved CNV data were estimated by GISTIC2 and thresholded, with -2, -1, 0, 1, 2 indicating two copy deletions, one copy deletion, no change, amplification and high amplification.^18^ Somatic mutation data was determined from the MC3 project, including non-silent mutation such as single nucleotide polymorphisms (SNPs) and insertion-deletions (INDELs).^19^ Binary mutation call was used as the omics data in our study.

### Methylation

We obtained β values from the Illumina HumanMethylation450 array, which served as our representation of DNA methylation at CpG sites.^20^ As the methylated probes *p* ∈ ℝ have different regulatory meanings across different locations in a gene, we grouped the CpG probes along a CDg into CpG island, non-CpG island, promoter, non-promoter, and enhancer according to the probeMap file based on hg38 reference genome.^21^ Each CDg has the probe vector *P^i^* = ( *p*_1_, … , *p_m_*) with length m. To create a uniform structure, we padded the lists with zero so that each CDg had the same maximum length, M, for a specific cancer type: 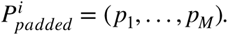

To transform the distribution of methylated β values into a compact, low-dimensional format while preserving biological meaning across distinct genomic regions, we employed an Autoencoder (AE) model. During the forward pass through the AE, input DNA methylation data underwent dimensionality reduction facilitated by three encoder layers. LeakyReLU activation functions introduced non-linearity to capture intricate patterns in the data. Additionally, we incorporated an optional masking mechanism into the AE to address variable-length methylation data. This mechanism facilitated the exclusion of padded zeros, ensuring that only relevant information contributed to the encoding process. The masking operation involved element-wise multiplication of the encoded representation by a binary mask, generated based on data presence or absence in the input. The output of this AE model is a compact, low-dimensional representation of DNA methylation data for each CDg *f_AE_* : *P_padded_* ∈ ℝ*^M^* → *P_enc_* ∈ ℝ*^n^*, where n is set to 1 in our study. The low-dimensional output was verified and visualized by t-SNE, based on the mean methylation values across all methylated probes in different regulatory regions.

### Determination of Cancer Drivers

DriverDBv3 (http://ngs.ym.edu.tw/driverdb), an integrative multi-omics database for cancer-related molecular intricacies, was used to extract CDgs for different cancer types.^6^ This invaluable resource employs well-established bioinformatics algorithms to discern CDgs. Additionally, DriverDBv3 extends its capabilities by elucidating the roles of CNV and methylation drivers, providing essential molecular features that contribute to a comprehensive understanding of ‘dysregulation’ events in cancer.

### Labels for binary classification tasks

To explore the ability of DriverOmicsNet to predict cancer-related features of clinical value, homologous recombination deficiency (HRD), cancer stemness, immune cluster, tumor stage and survival were utilized for binary classification.^22,23^ HRD, stemness (based on transcriptomic feature) and immune signature (IM) scores^24^ were obtained from analytic data type in the Xena browser using the TCGA-Pan-Cancer hub. For HRD and stemness scores, we established binary groupings based on their respective median values (Supplementary Fig. 1 and 2). The immune signature (IM) consisted of 68 signatures, and grouping within the IM was determined using hierarchical clustering performed with the *hclust* function in R (Supplementary Fig. 3). Additionally, we extracted tumor pathologic stage and overall survival data for each cancer type. For stage grouping, we classified tumors into early stage (stage 1 and 2) and late stage (stage 3 and 4), while survival labels were assigned based on median survival calculations.

### Overview of DriverOmicsNet

DriverOmicsNet is a comprehensive framework designed for addressing classification tasks involving multi-omics data. The workflow of DriverOmicsNet can be distilled into two key components (Fig. 1): (1) Network construction, which entails the generation of networks based on STRING PPI and correlation-based WGCNA. (2) Fusion of latent vectors for binary classification, where the final prediction is reached by concatenating the vector embeddings generated from the last layers of both GCN models. This integration of latent vectors from diverse sources enhances the framework’s capability to execute binary classification tasks effectively.

**Figure 1.**
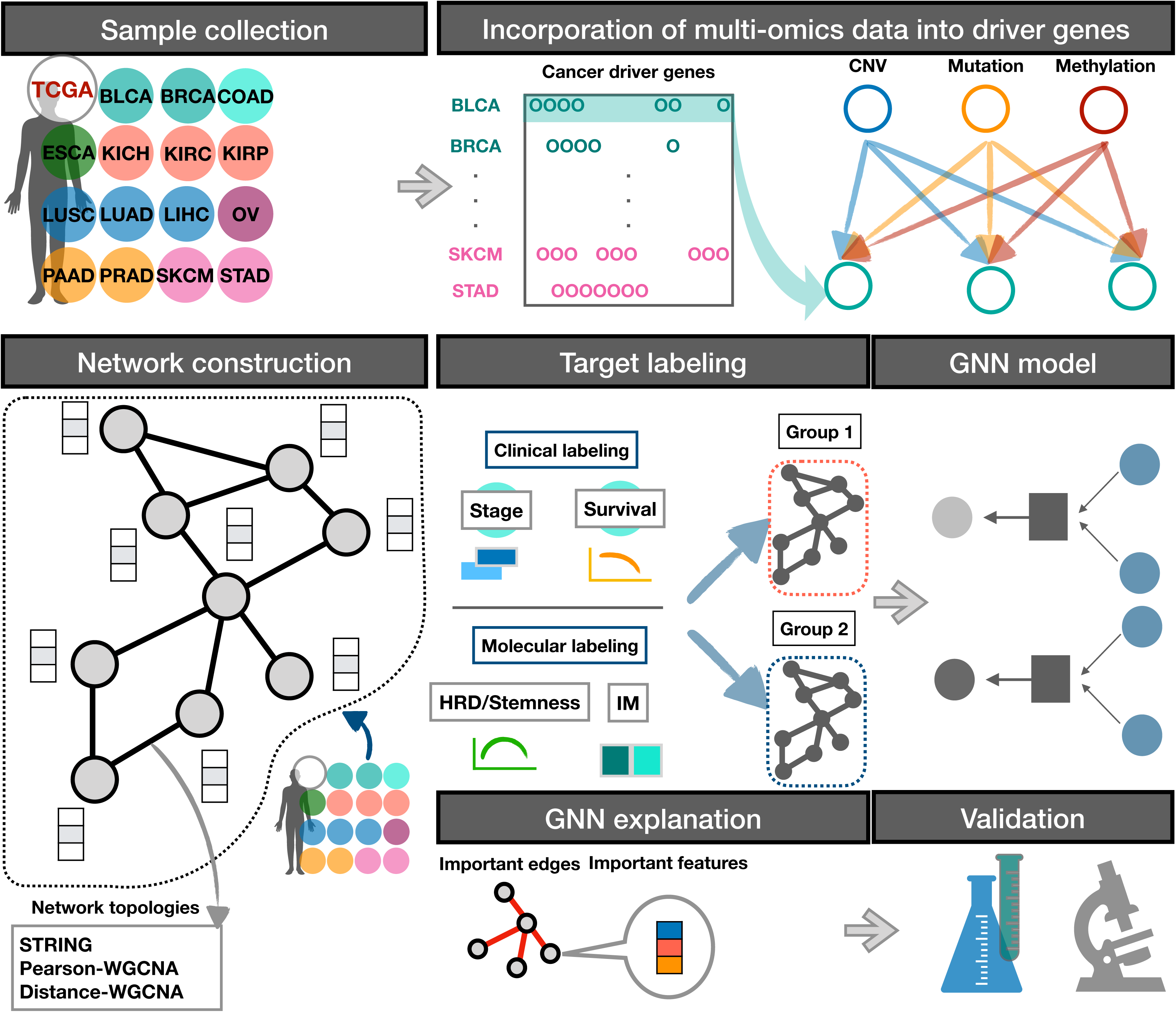
Overview of DriverOmicsNet.

### GCN for DriverOmicsNet

The goal of DriverOmicsNet is to take advantage of GCN and explore the key graphlets that is crucial for label classification. First, STRING PPI graph *g*_*p*(*v*,*e*)_ and correlation-based WGCNA graph *g*_w(*v*,*e*)_ were constructed for each cancer type, where *v* and *e* represented node set and edge set, respectively. STRING is a well-established database for experimentally-verified interactions among query proteins, therefore providing a rich foundation for our analysis.^25^ For *g*_*p*(*v*,*e*)_, undirected edges connecting each CDg were defined based on a combined score > 0.4. For constructing *g*_w(*v*,*e*)_, we first obtained a similarity co-expression matrix among CDgs. In addition to traditional Pearson’s correlation, we used Distance correlation (*dcor* package in python) to explore its ability of capturing biologically meaningful networks.^26^ Hub genes for each network were defined as the genes with the highest node degrees. Next, adjacency matrices were obtained by using the soft-thresholding power, which was selected to approximately fit a scale-free topology. Subsequently, topological overlap matrices *TOM_w_*_,*p*_ and *TOM*_*w*,*d*_ were generated. To enhance the comparability with *g*_*p*(*v*,*e*)_, thresholding was applied for each cancer type to approximate the edge set number of 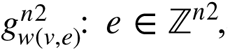 with 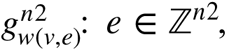 so that *n*1 ≈ *n* 2.

In order to integrate multi-omics data in the processes of message passing and aggregation for GCN, the graphs 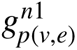 and 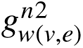 were converted to data structures acceptable for training a GCN model, which is a node feature matrix *X* ∈ ℝ*^nxk^* where is the number of CDgs and *k* is the feature size (size=8 gene expression, CNV, mutation, CpG island, non-CpG island, promoter, non-promoter, and enhancer), and a data structure called edge index *E* = [(*u*_1_, *v*_1_), (*u*_2_, *v*_2_), … (*u_m_*, *v_m_*)] ∈ ℤ^2^*^xm^* where (*u*, *v*) represents a pair of node indices connected by an edge and denotes the total number of edges in an undirected graph. For *g*_w(*v*,*e*)_, edge attribute for each edge is given as an additional parameter, which is the value from *TOM_w_*_,*p*_ and *TOM*_*w*,*d*_.

Specifically, for 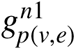 and 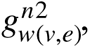 the GCN models can be denoted as 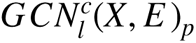 and 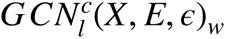 where *l* ∈ ℤ^5^ is the target label of interest and *c* ∈ ℤ^15^ is the cancer type. Generally, each model has the same structure, with four GCN layers and three fully connected layers (MLPs). Each GCN layer is activated by ReLU, which is followed by TopKPooling to reduce the number of nodes while retaining the most informative ones. The pooled nodes are then globally pooled to obtain a graph-level representation, resulting in four layers *L* = (*L*_1_, *L*_2_, *L*_3_, *L*_4_) ∈ ℝ*^kxh^* where is is the hidden dimension during training. The four layers are concatenated to create a comprehensive graph-level representation *L*\* : *concat* (*L*_1_, *L*_2_, *L*_3_, *L*_4_) ∈ ℝ*^kxh^*, which is followed by the three MLPs. The output of the last linear layer is passed through a softmax function to obtain class probabilities for classification. Configuration of the GCN structure is shown in Supplementary Figure 4.

### DriverOmicsNet for binary classification

We randomly split the total samples of each cancer type into training and testing sets with a 8:2 ratio. We fine-tuned the hyperparameters (batch size, hidden dimension, learning rate and weight decay rate) for 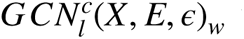 and 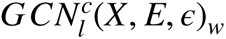 with HyperOPT and Ray.^27^ Maximal number of epochs of 500 with early stopping was used with the patience of 10. Initially, 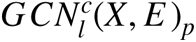 and 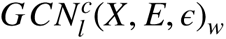 were trained in an end-to-end fashion. To establish a final model that is able to capture local network structure from STRING PPI and WGCNA networks, *L*\* from each model 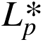 and 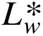 are concatenated, forming 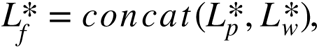 which is followed by two MLPs to make final prediction (Supplementary Fig 5)

### Model performance metrics

To thoroughly gauge the effectiveness and robustness of our GCN models, we conducted a comprehensive evaluation employing both a fivefold cross-validation and independent testing procedures. The evaluation outcomes were summarized based on key performance metrics, including accuracy, F1-Score, Matthews Correlation Coefficient (MCC), and the area under the receiver operating characteristic curve (AUC-ROC).

### Identification of key CDgs for binary classification

We initiated the GNN Explainer with our GCN model architecture and conducted an iterative training process for 100 epochs. During each epoch, we determined the node and edge importance for each input graph in our dataset. The node importance was quantified through a node feature mask, and edge importance was captured using an edge mask. These masks were used to highlight the most critical nodes and edges driving the GNN’s predictions. Next, we applied a similar procedure for the distance network. The cumulative importance scores of nodes and edges were computed throughout the datasets, enabling us to identify the top 100 important nodes and edges based on their significance scores. We established a threshold using the 100th highest score, ensuring that only the most influential nodes and edges were retained for visualization. Subsequently, we created subgraphs composed of these critical elements, visualizing them to gain insights into their structural and functional importance within the networks.

## Results

### Multi-omics features of driver genes in cancer

Among the 15 cancer types, DriverDBv3 has identified a total of 2620 CDgs. These genes play pivotal roles in driving cancer progression and are integral components of the TCGA cohorts. The intricate inter-correlations among various omics layers for these CDgs are summarized in Figure 2a, as detailed in the database’s driver summary table (http://driverdb.bioinfomics.org). Furthermore, performing ORA on these drivers has unveiled cancer signaling as the most enriched biological term, underscoring the substantial involvement of these CDgs in cancer-related processes (Fig. 2b). Furthermore, when assessing the GS of each CDg, a robust correlation emerged between mRNA expression and the tumor trait (Fig. 2c). It is noteworthy that a notable subset of CDgs exhibited negative GS values across different cohorts. Among these, some represent oncogenes frequently characterized by their overexpression in various cancer types, such as *PDGFRA*, *PIK3R1*, and *BRAF* (Fig. 2d).^28–30^ Conversely, *TP53* consistently showed positive GS values, emerging as the top CDg across all cancer cohorts. This finding reinforces the established evidence that *TP53* is a pivotal driver gene in cancer.^31^ In addition to gene expression, there is a varying distribution of CNV and mutational status among the CDgs for each cancer type. By pooling the number of CDg mutations across all study samples, *TP53* emerges as the most commonly mutated CDg for all cancer types, followed by *PIK3CA,* which is consistently observed in cancer patients from different populations (Supplementary Fig. 6).^32^ Taken together, the processed data extracted from TCGA represents a broad spectrum of cancer patients, and the diverse distribution of each omics data type helps capture the varying landscapes of multi-omics in cancer, allowing GNN models to generalize better to different scenarios and make more accurate predictions or classifications.

**Figure 2.**
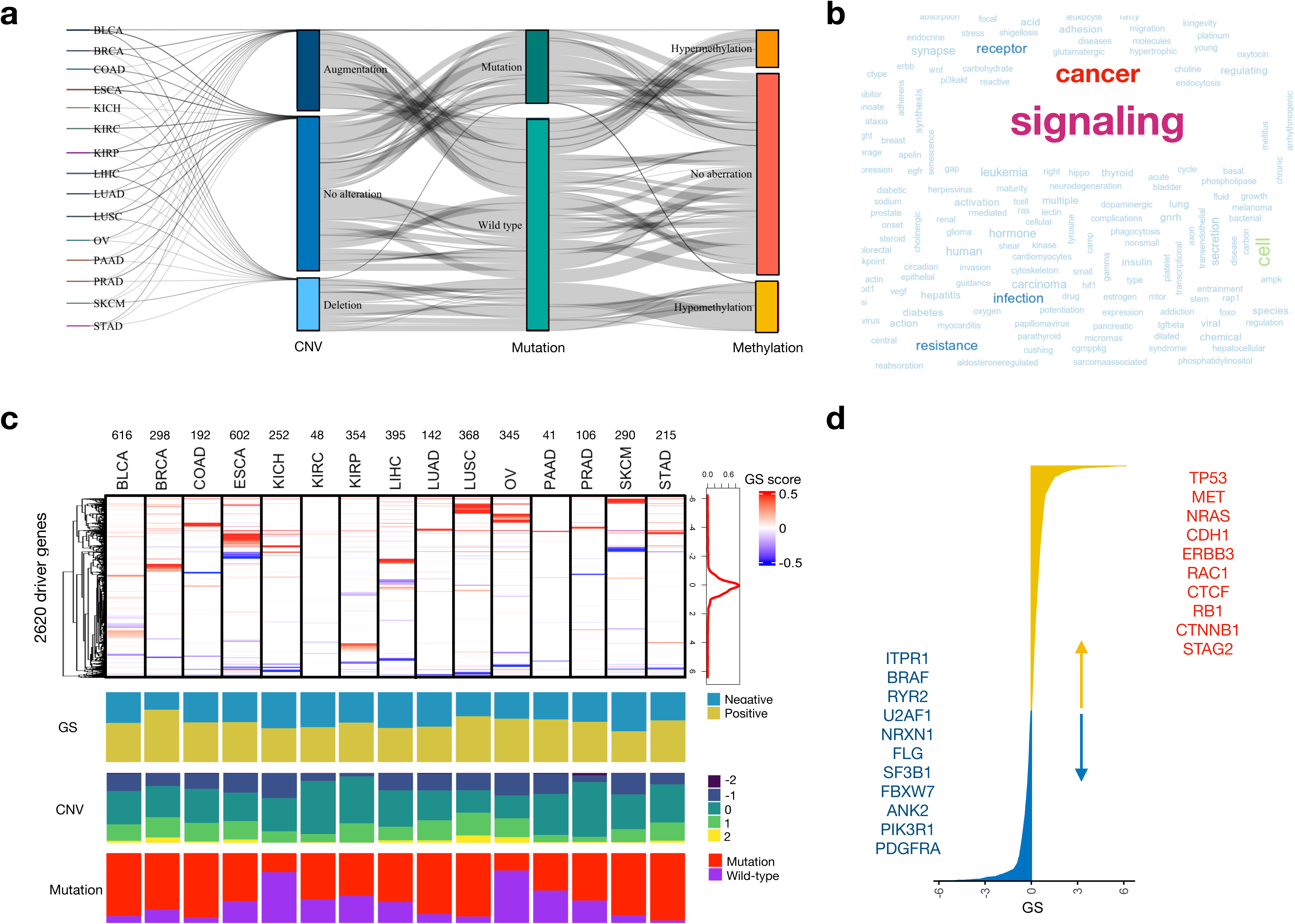
Characterization of the cancer driver genes (CDgs) used in the study for each cancer type. **a**. Sankey diagram showing the associations among copy number variations, mutations and abnormal methylations for CDgs. Nodes in the graph with more connections suggest a greater influence **b**. Enriched gene ontology terms for all CDgs across all cancer types. c. Distributions of GS scores derived from WGCNA, CNVs and somatic mutations across all cancer types. d. Cumulative GS scores for all CDgs across all cancer types. WGCNA=weighted gene co-expression network analysis. GS=gene significance. CNV=copy number variation.

### Encoding of DNA methylation data

The distribution of β values for each genomic region is summarized for every cancer type in Supplementary Fig. 7, showing varying methylation patterns specific for each region. The AE model reduced the dimensionality of the input data through a series of encoder layers, incorporating LeakyReLU activation functions to capture intricate patterns within the DNA methylation profiles (Fig. 3a-b). We observed distinct clusters of DNA methylation status in many cancer types (Fig. 3c and Supplementary Fig. 8), signifying the effectiveness of the AE model in capturing underlying methylation patterns.

**Figure 3.**
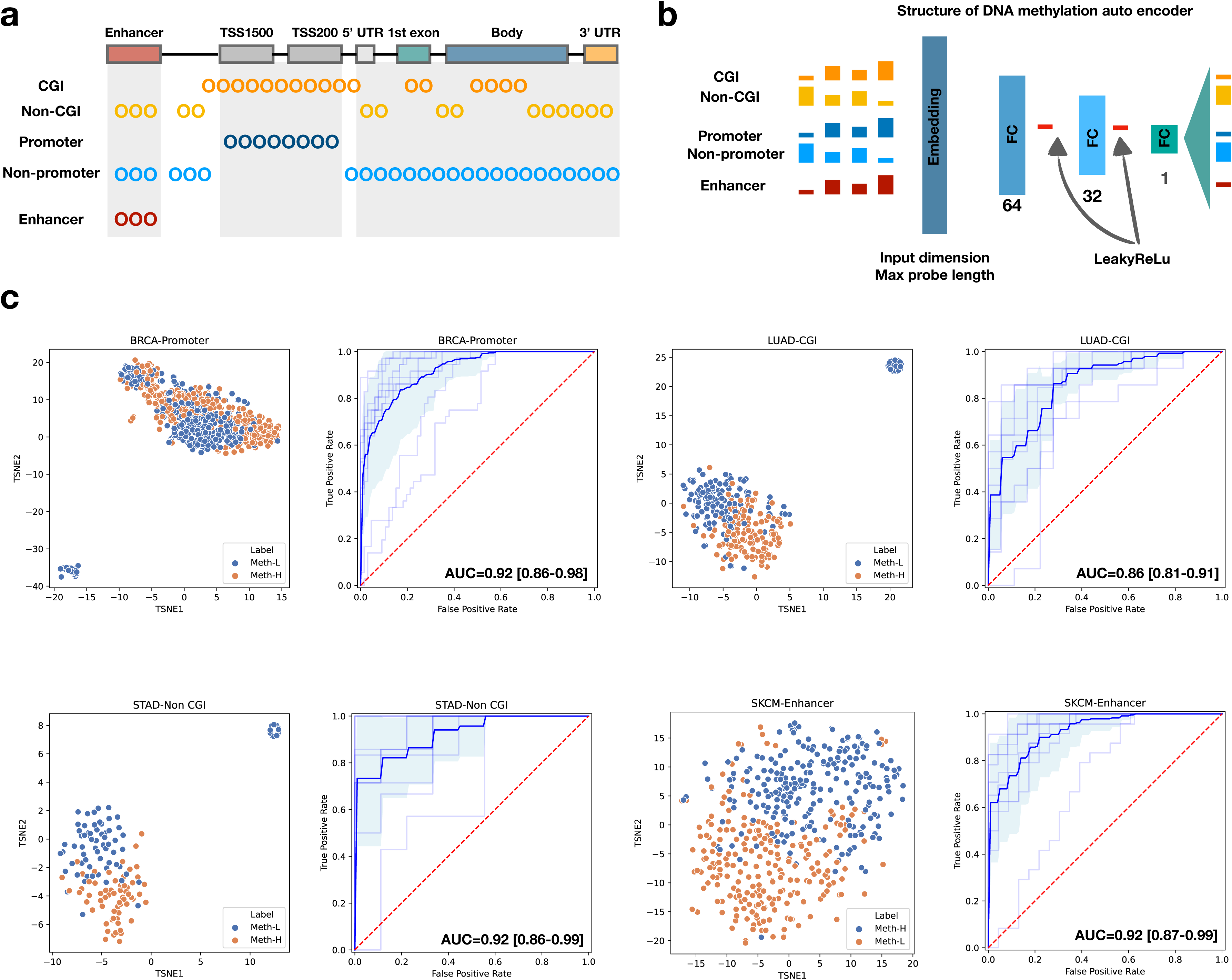
Encoding of methylation for multi-omics integration. **a**. Different regulatory regions of methylation outlined in HumanMethylation450 array. **b**. Configuration of DNA methylation autoencoder (AE). **c**. t-SNE plots and area under curve plots showing distribution of decoded outputs from AE and their capabilities of predicting mean methylation statuses in BRCA, LUAD, STAD and SKCM across different regulatory regions. BRCA= Breast invasive carcinoma. LUAD= Lung adenocarcinoma. STAD= Stomach adenocarcinoma. SKCM= Skin Cutaneous Melanoma.

Furthermore, we leveraged a MLP model to predict the methylation status based on the converted data. Encouragingly, our predictions achieved remarkable accuracy with only one-dimensional data as the input for MLP, with an AUC of 0.92 (95% CI: 0.86-0.98) for promoter region in breast invasive carcinoma (BRCA), 0.86 (95% CI: 0.81-0.91) for CpG island in lung adenocarcinoma (LUAD), 0.92 (95% CI: 0.86-0.99) for non-CpG island in stomach adenocarcinoma (STAD), and 0.92 (95% CI: 0.87-0.99) for enhance in cutaneous melanoma (SKCM), underscoring the potential of this approach in simplifying and stratifying DNA methylation patterns (Fig. 3c).

### Predicting outcomes based on separate network structure

The number of undirected edges derived from STRING PPI varies across different cancer types, closely mirroring the respective input gene numbers (Supplementary Fig. 9).

Supplementary Fig. 10 illustrates the thresholds of correlation coefficients used to ensure the comparability of PPI and CDg co-expression modules. In the case of 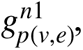 the hub genes remain consistent across all cancer cohorts, with *TP53* exclusively identified as the hub gene (Supplementary Table 1). However, for 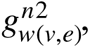 there is a striking divergence in hub genes across various cancer types. Most cancer types exhibit shared hub genes between the Pearson- and Distance-WGCNA networks, except for a few exceptions, including esophageal carcinoma (ESCA), liver hepatocellular carcinoma (LIHC), ovarian serous cystadenocarcinoma (OV), and STAD.

Based on the network structures, a total of 225 GCN models were meticulously evaluated. Notably, we observed that the WGCNA-based models consistently outperformed the STRING-based models across all five target labels (Fig. 4a). Moreover, when comparing the performance between Distance and Pearson correlation-based models, we found a slight but consistent improvement in model accuracy with the Distance WGCNA approach. Therefore, we selected STRING- and Distance WGCNA-based models for combination. Additional 75 models were trained, and our results unveiled a substantial improvement in model training and testing accuracy when applying the concatenated model that fuses the strengths of both STRING PPI and WGCNA networks (Supplementary Fig. 11). This combined approach demonstrated remarkable enhancements of testing accuracy for the five label classes, ranging from average 0.1% for HRD to a substantial 15.2% for survival across various cancer types, as compared with Distance WGCNA- and STRING-based models, respectively (Fig. 4b).

**Figure 4.**
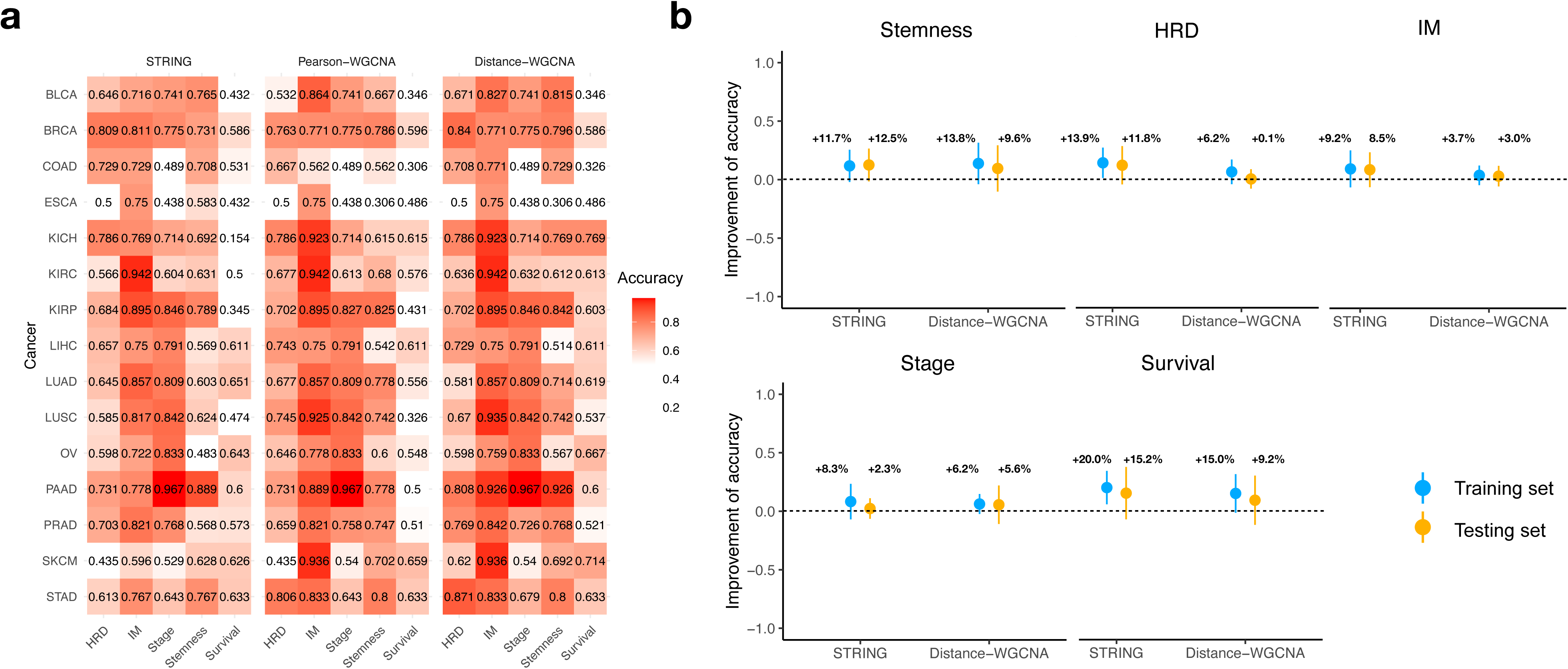
Model accuracies of graph convolutional networks (GCN). **a**. Model testing accuracies based on network structures of STRING, Pearson correlation-based WGCNA, and Distance correlation-based WGCNA across five target features. Color bar indicates the testing accuracy. **b**. Improvement of accuracies observed in DriverOmicsNet. Improvements of accuracy are displayed as mean±standard deviation for training and testing sets. GCN models built from STRING and Distance correlation-based WGCNA networks are set as the baseline. WGCNA=weighted gene co-expression network analysis.

### The top 30 combined models

To assess the potential of our combined models in predicting specific labels, we focused on the top 30 models, which represent the upper 40% in terms of predictive performance among all combined models. These models were primarily trained for predicting IM, with a secondary focus on stemness and stage prediction (Supplementary Fig. 12). Notably, there were no models dedicated to predicting survival among the top-performing ones, suggesting limited predictive value in this context. Among the top ten models, seven were designed for predicting IM clusters, achieving testing accuracy ranging from 89.47% for kidney renal papillary cell carcinoma (KIRP) to 95.15% for kidney renal clear cell carcinoma (KIRC). Subsequently, we selected models with the highest testing accuracy for HRD, stemness, IM, and stage predictions to evaluate their performance. The selected models corresponded to STAD (for HRD prediction), KIRP (for stemness and stage prediction), and SKCM (for IM prediction) (Fig. 5). Among these models, the one predicting HRD status for STAD exhibited the best performance metrics (Precision=0.9307 ± 0.0108; F1 score=0.9102 ± 0.0938; MCC=0.8368 ± 0.1540, and AUC=0.9604 ± 0.0642), followed by the model predicting IM clusters for SKCM (Precision=0.9091 ± 0.0656; F1 score=0.8749 ± 0.1078; MCC=0.7745 ± 0.1837, and AUC=0.9195 ± 0.0960). The model for predicting stage in KIRP showed the poorest performance among the four label classes. The label ratio of IM for SKCM is 1.05, and the ratio of stage for KIRP is 2.33 (Supplementary table 2). We also explored additional two models for predicting stage with a relatively small label ratio. However, it’s important to note that the additional models still exhibited comparatively poorer performance (data not shown).

**Figure 5.**
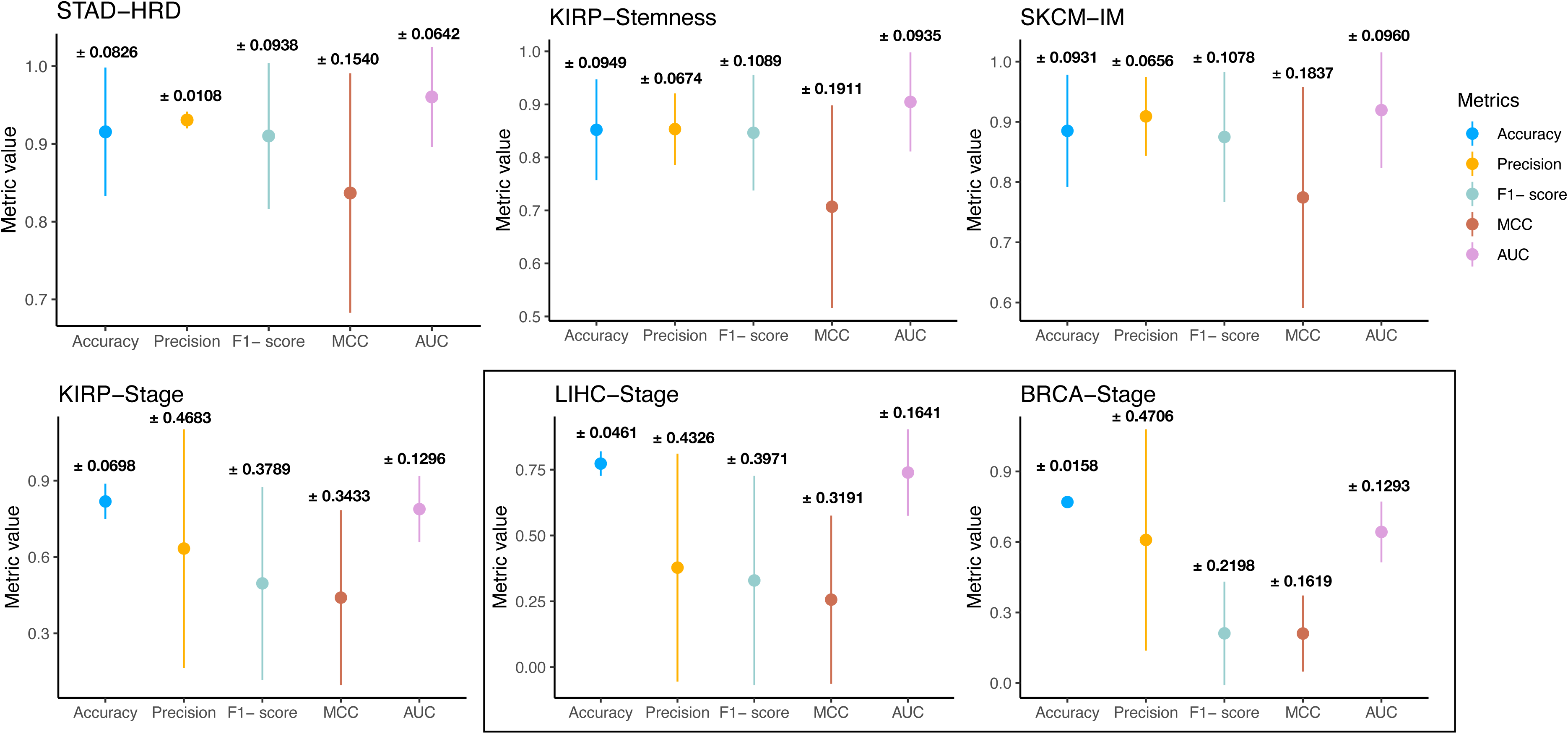
Model performance for selected models with the highest testing accuracy for HRD, stemness, IM, and stage predictions. Additional two models are demonstrated for stage. STAD= Stomach adenocarcinoma. KIRP= Kidney renal papillary cell carcinoma. SKCM= Skin Cutaneous Melanoma. LIHC= Liver hepatocellular carcinoma. BRCA= Breast invasive carcinoma. MCC= Matthews Correlation Coefficient. AUC=area under curve.

### Omics importance and graphlet identification for molecular features

To explore the contribution of each omics feature to graph prediction, we selected models with balanced labels (label ratio < 3) to ensure robust performance while minimizing bias. Consequently, we retained a total of 17 models. The cumulative node feature importance and subgraphs with overlapping CDgs derived from STRING and Distance WGCNA models are presented in Supplementary Fig. 13. Overall, gene expression emerged as the most significant feature for graph prediction, followed by CNV in both graph structures. In contrast, mutation status exhibited the lowest importance value. Methylation data showed substantial variation across different cancer types and graph structures. This variability also extended to the subgraphs, reflecting diverse degrees of connectivity among nodes. Taken together, these findings underscore the paramount importance of gene expression and CNV in predicting graph structures, while highlighting the need to consider cancer-type-specific differences when incorporating methylation data.

## Discussion

Our study demonstrates the effectiveness of using a GCN to integrate multi-omics data for predicting cancer-related features, especially HRD, cancer stemness, and immune clusters. The robustness of DriverOmicsNet in integrating data from STRING PPI networks and correlation-based WGCNA networks underscores its potential in precision oncology. Our study also introduces a novel approach to processing and generating multi-omics data, with a particular focus on the innovative handling of DNA methylation data. Unlike traditional methods that treat different methylation sites uniformly, we have developed a sophisticated AE model that reduces the dimensionality of methylation data while preserving biologically significant patterns. Consequently, our method not only simplifies and stratifies DNA methylation patterns but also significantly improves the predictive power and biological relevance of the generated data, thereby advancing the field of precision oncology.

While the STRING PPI network alone is comprehensive and frequently updated, it has several notable limitations.^33^ The coverage of experimentally verified interactions for specific conditions, such as Alzheimer’s disease, is relatively low compared to other databases, indicating potential gaps in STRING’s dataset for certain disease-specific interactions.^25^ Additionally, the classification of interactions can vary, with interactions considered experimentally verified in STRING not always categorized similarly in other integrated databases like hPRINT.^34^ This inconsistency can lead to confusion and misinterpretation of the data. Nevertheless, STRING still offers several significant advantages. Its extensive coverage includes over 5090 organisms and 24.6 million proteins, providing a broad and diverse dataset for researchers.^33^ STRING’s integration of multiple data sources—experimental data, computational predictions, and text mining— ensures a comprehensive view of protein interactions. Constructing PPI networks based on WGCNA co-expression networks offers unique and additional advantages. WGCNA constructs co-expression networks that reflect condition-specific gene expression patterns, allowing for the identification of interactions relevant to specific tissues, developmental stages, or disease states.^35^ It captures dynamic changes in gene expression, identifying interactions that may only occur under certain conditions or in response to specific stimuli, which is often missed in STRING’s static compilations.^36^ Furthermore, WGCNA directly incorporates gene expression data, making it easier to correlate network properties with expression levels and understand the regulatory mechanisms underlying gene co-expression. Additionally, WGCNA provides quantitative measures of gene connectivity within modules, which can be used to identify hub genes that play central roles in network function. This quantitative approach, combined with the focus on co-expression, reduces the inclusion of false positive interactions that can arise from STRING’s aggregation of heterogeneous data sources. While STRING’s comprehensive integration approach is valuable, it can sometimes introduce spurious interactions that are not relevant in specific biological contexts.

Another significant advantage of using WGCNA is the flexibility in choosing the method for calculating co-expression. Specifically, using distance-based co-expression over Pearson correlation in the WGCNA algorithm provides several benefits. Distance-based measures can capture non-linear relationships between genes that Pearson correlation might miss.^26^ This is particularly useful in complex biological systems where gene interactions are not strictly linear. Distance-based methods also tend to be more robust to outliers, providing a more stable and reliable network structure. Moreover, they can better handle variations in gene expression levels across different conditions, leading to more accurate identification of gene modules and interactions. This flexibility enhances the relevance and accuracy of the network, particularly in diverse and heterogeneous datasets such as those encountered in cancer research.

The utilization of GCNs in our study offers several advantages. GCNs excel at learning from graph-structured data, enabling the integration of multi-omics features into a cohesive network representation. This is particularly beneficial when combining STRING PPI and WGCNA networks, as GCNs can effectively capture the complex relationships between genes and their interactions.^37^ Unlike the MOGONET framework by Wang et al., which used separate GNNs for each omics data type, our unified approach integrates all data types into a single model, providing a holistic view of the molecular landscape.^11^ By leveraging both the static, experimentally verified interactions from STRING and the dynamic, condition-specific interactions from WGCNA, GCNs enhance the predictive power of our models and facilitate the identification of key driver genes and their interactions. This integrative approach advances our understanding of cancer biology and informs precision oncology strategies.

Our results underscore the critical role of CDgs and highlight gene expression as the most pivotal molecular feature across various cancer types and clinical outcomes. The prominence of gene expression in our GCN models is due to several factors. Firstly, gene expression directly reflects the functional state of cells, offering immediate insights into the biological processes and pathways involved in cancer progression. This makes it a highly informative marker for distinguishing between different cancer phenotypes and predicting clinical outcomes. Additionally, the dynamic nature of gene expression allows for the capture of condition-specific changes, essential for understanding tumor adaptation and response to their microenvironment and treatment. Furthermore, the integration of multi-omics data through the WGCNA framework amplifies the importance of gene expression by correlating it with other genomic alterations, such as CNVs and mutations, providing a comprehensive view of the molecular underpinnings of cancer. This integrative approach reduces the likelihood of identifying false-positive driver genes, enhancing the accuracy of our findings.^38^

Despite the promising results of our study, several limitations should be acknowledged. Firstly, the integration of multi-omics data, while comprehensive, is dependent on the quality and completeness of the available datasets. Any gaps or biases in the data could impact the accuracy and generalizability of our models. Secondly, while our approach successfully combines static and dynamic interaction networks, it does not fully account for temporal changes in gene expression and protein interactions over time, which are crucial in understanding cancer progression and treatment response. Moreover, the complexity of GCN models requires substantial computational resources, potentially limiting their accessibility for broader clinical application. Finally, while our models have shown robust predictive power across multiple cancer types, their performance in rare cancers or in datasets with limited sample sizes remains to be fully validated. Future studies should aim to address these limitations by incorporating more diverse datasets, improving data annotation consistency, and exploring methods to reduce computational demands.

## Conclusion

In conclusion, the integration of STRING and WGCNA networks within DriverOmicsNet leverages the strengths of both approaches, enhancing the robustness and relevance of the resulting PPI networks for precision oncology applications. This dual approach provides a more nuanced and comprehensive understanding of cancer biology, facilitating the development of more effective predictive models and therapeutic strategies. Furthermore, the use of distance-based co-expression in WGCNA adds an additional layer of robustness and accuracy, making it a powerful tool for constructing biologically meaningful networks.

