## Supplementary material for "DriverOmicsNet: An Integrated Graph Convolutional Network for Multi-Omics Exploration of Cancer Driver Genes"

**Supplementary Table 1.** Hub genes identified in each network structure.

| Cancer | STRING | Pearson-based WGCNA | Distance correlation-based WGCNA |
|---|---|---|---|
| BLCA | *TP53* | *TRAK1* | *TRAK1* |
| BRCA | *TP53* | *DSCC1* | *DSCC1* |
| COAD | *TP53* | *MOCS3* | *MOCS3* |
| ESCA | *TP53* | *TP63* | *HNF4A* |
| KICH | *TP53* | *STAG2* | *STAG2* |
| KIRC | *TP53* | *HERC1* | *HERC1* |
| KIRP | *TP53* | *CFAP69* | *CFAP69* |
| LIHC | *TP53* | *VPS28* | *EXOSC4, VPS28* |
| LUAD | *TP53* | *MRPS28* | *MRPS28* |
| LUSC | *TP53* | *ZNF639* | *ZNF639* |
| OV | *TP53* | *FBN3* | *DNMT3A, THEM6, GSDMC* |
| PAAD | *TP53* | *BCL2* | *BCL2* |
| PRAD | *TP53* | *IQCA1* | *IQCA1* |
| SKCM | *TP53* | *ROCK1, STAG2* | *ROCK1, STAG2* |
| STAD | *TP53* | *SYNE1, ITPR1* | *SYNE1* |

WGCNA=weighted gene co-expression network analysis; BLCA= Bladder Urothelial Carcinoma; BRCA= Breast invasive carcinoma; COAD= Colon adenocarcinoma; ESCA= Esophageal carcinoma; KICH= Kidney Chromophobe; KIRC= Kidney renal clear cell carcinoma; KIRP= Kidney renal papillary cell carcinoma; LIHC= Liver hepatocellular carcinoma; LUAD= Lung adenocarcinoma; LUSC= Lung squamous cell carcinoma; OV= Ovarian serous cystadenocarcinoma; PAAD= Pancreatic adenocarcinoma; PRAD= Prostate adenocarcinoma; SKCM= Skin Cutaneous Melanoma; STAD= Stomach adenocarcinoma

Supplementary Table 2. Label ratios for different features

| Cancer | HRD | Stemness | IM cluster | Stage | Survival |
|---|---|---|---|---|---|
| BLCA | 1.06 | 1.00 | 1.40 | 2.13 | 1.01 |
| BRCA | 1.07 | 1.00 | 2.49 | 3.07 | 1.01 |
| COAD | 1.11 | 1.02 | 1.16 | 1.33 | 1.02 |
| ESCA | 1.12 | 1.02 | 3.5 | 1.59 | 1.04 |
| KICH | 1.58 | 1.06 | 1.87 | 2.37 | 1.09 |
| KIRC | 1.51 | 1.00 | 17.3 | 1.42 | 1.00 |
| KIRP | 1.20 | 1.01 | 10.36 | 2.33 | 1.01 |
| LIHC | 1.01 | 1.01 | 2.92 | 2.81 | 1.01 |
| LUAD | 1.01 | 1.01 | 5.08 | 3.87 | 1.03 |
| LUSC | 1.06 | 1.00 | 4 | 3.87 | 1.03 |
| OV | 1.06 | 1.01 | 3.68 | 10.8 | 1.02 |
| PAAD | 1.03 | 1.01 | 2.65 | 14 | 1.02 |
| PRAD | 1.01 | 1.01 | 5.90 | 1.54 | 1.00 |
| SKCM | 1.01 | 1.00 | 1.05 | 1.21 | 1.07 |
| STAD | 1.03 | 1.01 | 1.13 | 1.13 | 1.05 |

HRD=homologous recombinant deficiency; IM: immune; BLCA= Bladder Urothelial Carcinoma; BRCA= Breast invasive carcinoma; COAD= Colon adenocarcinoma; ESCA= Esophageal carcinoma; KICH= Kidney Chromophobe; KIRC= Kidney renal clear cell carcinoma; KIRP= Kidney renal papillary cell carcinoma; LIHC= Liver hepatocellular carcinoma; LUAD= Lung adenocarcinoma; LUSC= Lung squamous cell carcinoma; OV= Ovarian serous cystadenocarcinoma; PAAD= Pancreatic adenocarcinoma; PRAD= Prostate adenocarcinoma; SKCM= Skin Cutaneous Melanoma; STAD= Stomach adenocarcinoma

**Supplementary Fig. 1.** Heatmaps showing hierarchical clusters of 68 immune signature scores across all cancer types.

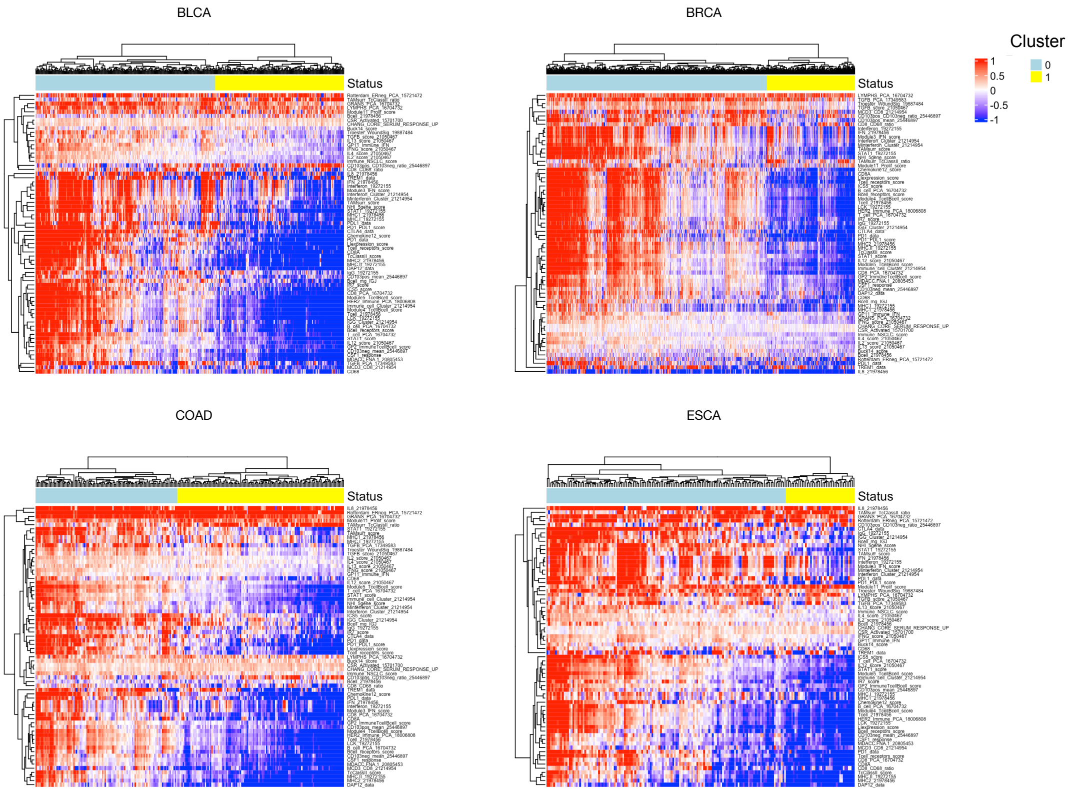

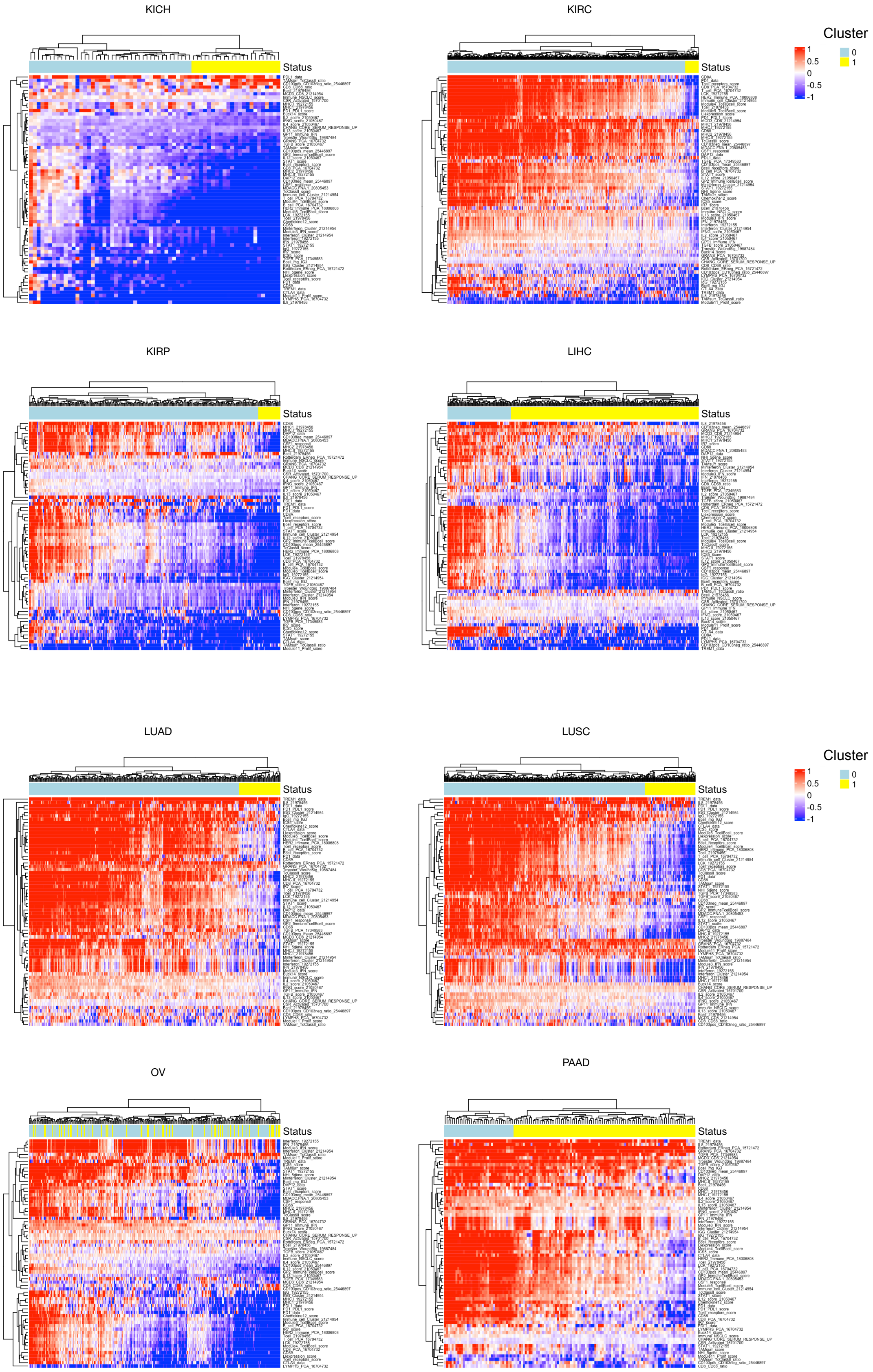
KICH
KIRC
KIRP
LIHC
LUAD
LUSC
OV
PAAD
Status
Cluster
1
0.5
0
-0.5
-1
0
1

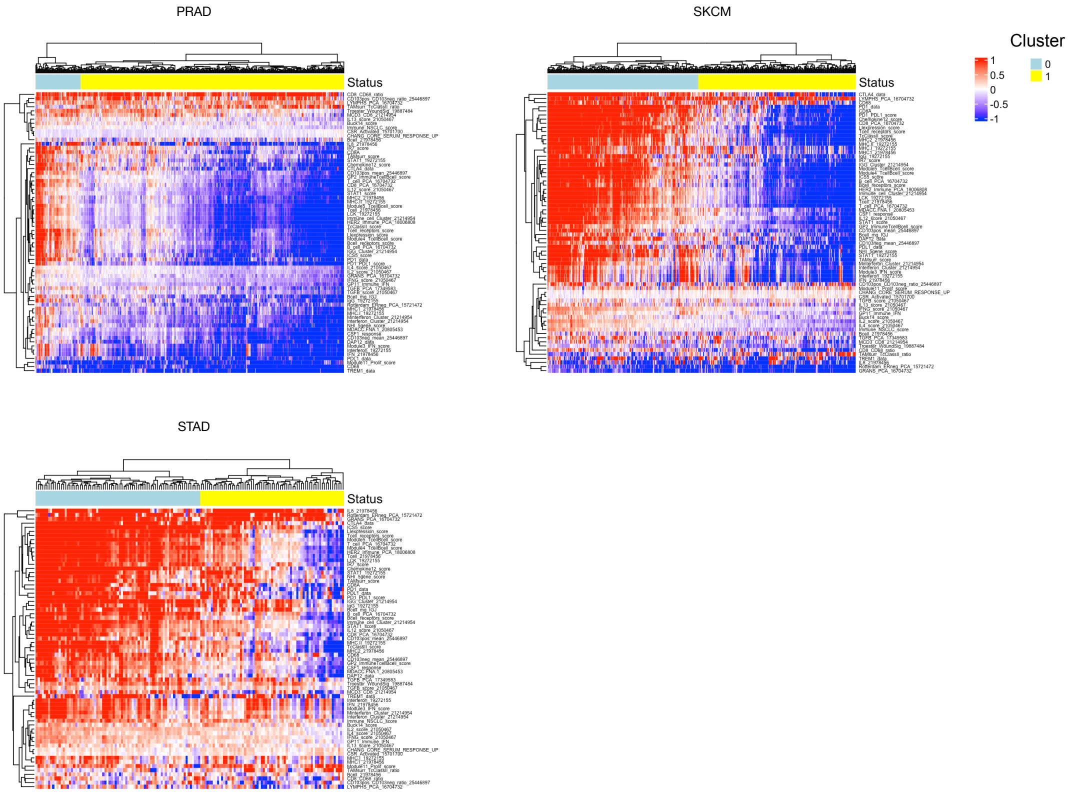
PRAD
SKCM
STAD
Status
Cluster
1
0.5
0
-0.5
-1
0
1

**Supplementary Fig. 2A.** Distribution of HRD scores across all cancer types.

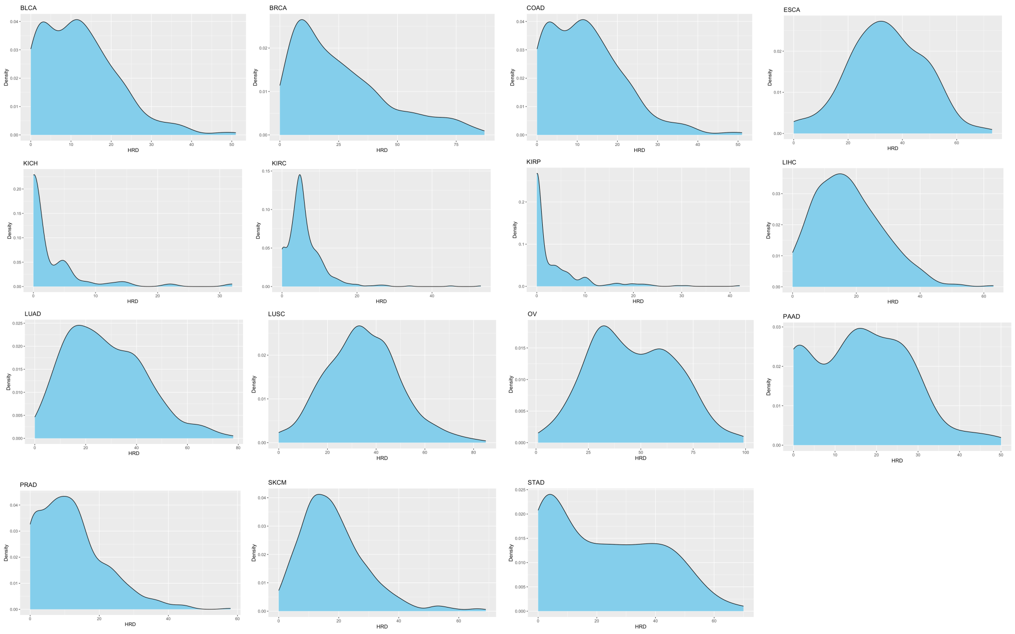


**Supplementary Fig. 2B.** Median HRD scores across all cancer types.

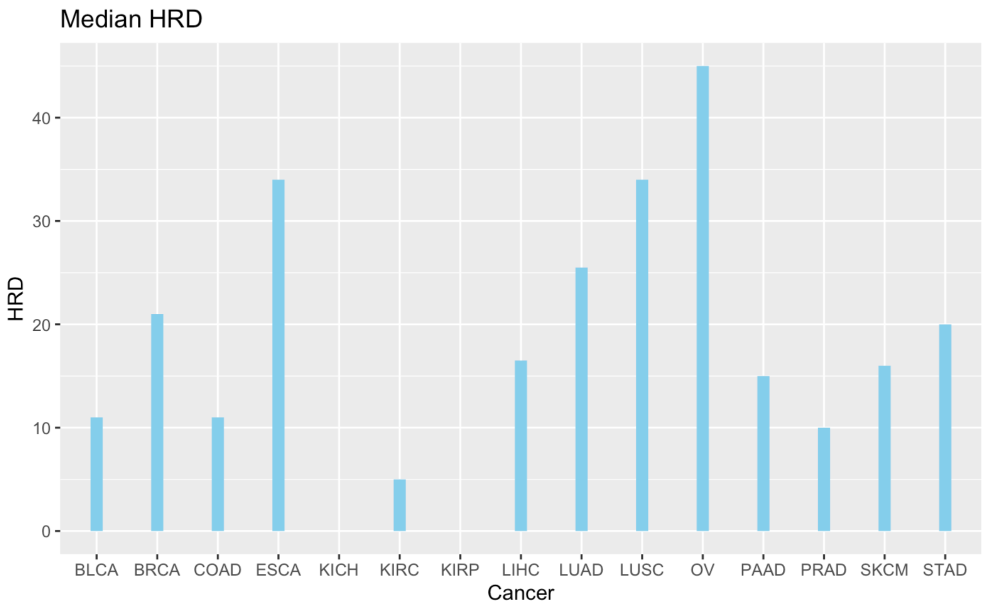

**Supplementary Fig. 3A.** Distribution of Stemness scores across all cancer types.

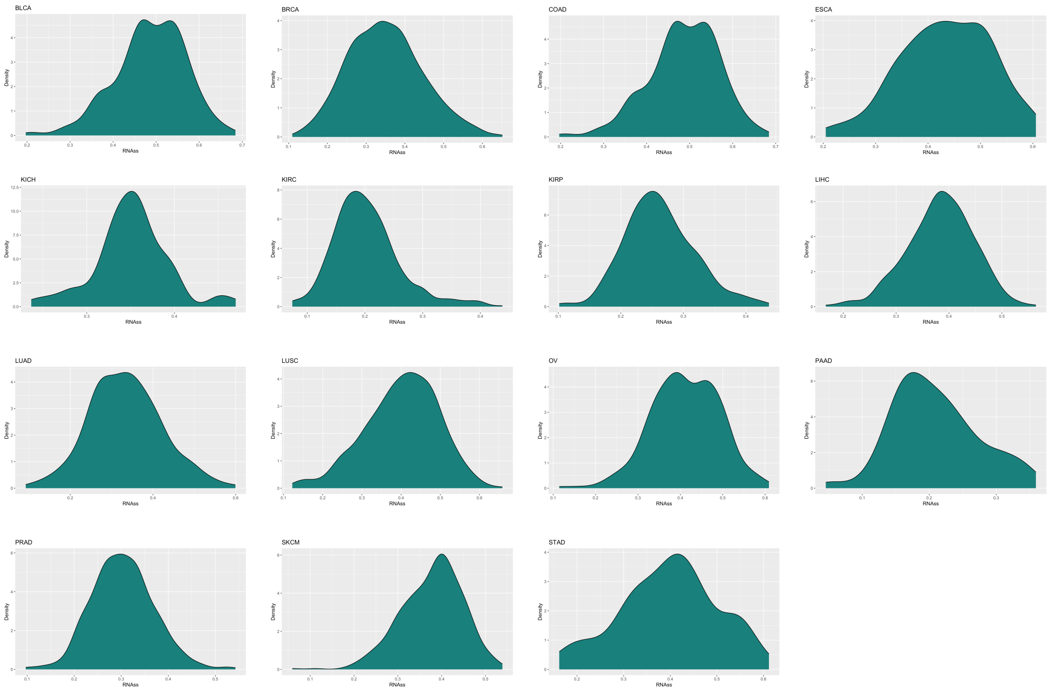


**Supplementary Fig. 3B.** Median Stemness scores across all cancer types.

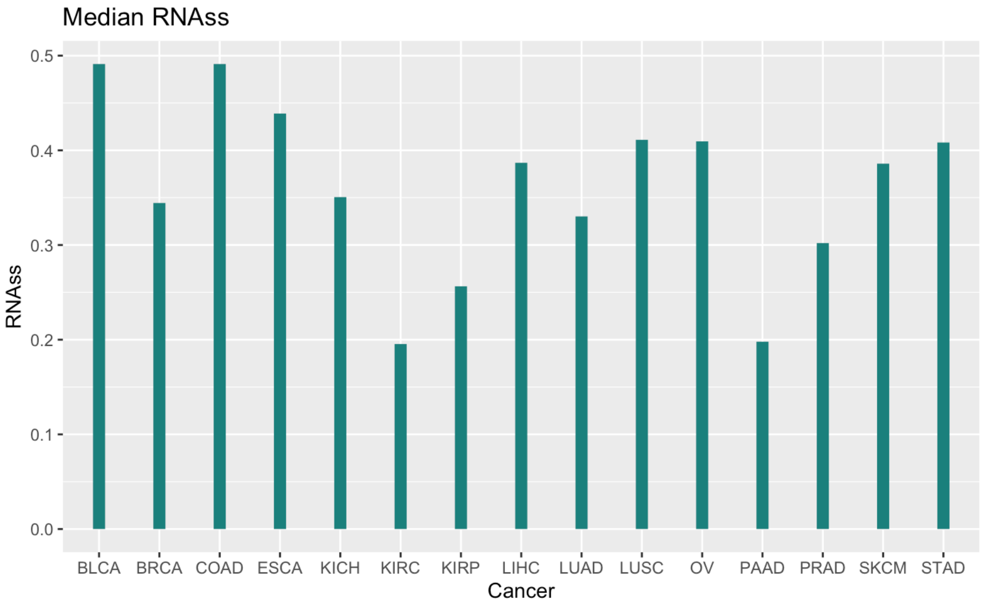

**Supplementary Fig. 4.** Configuration of GCN structure.

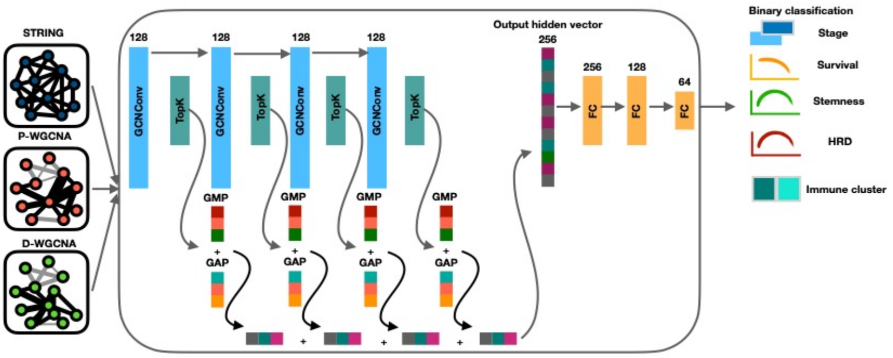


**Supplementary Fig. 5.** Configuration of the combined GCN model.

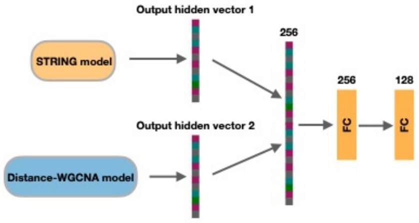

**Supplementary Fig. 6.** Cumulative counts of total somatic mutations across all cancer types for the top 20 cancer driver genes.

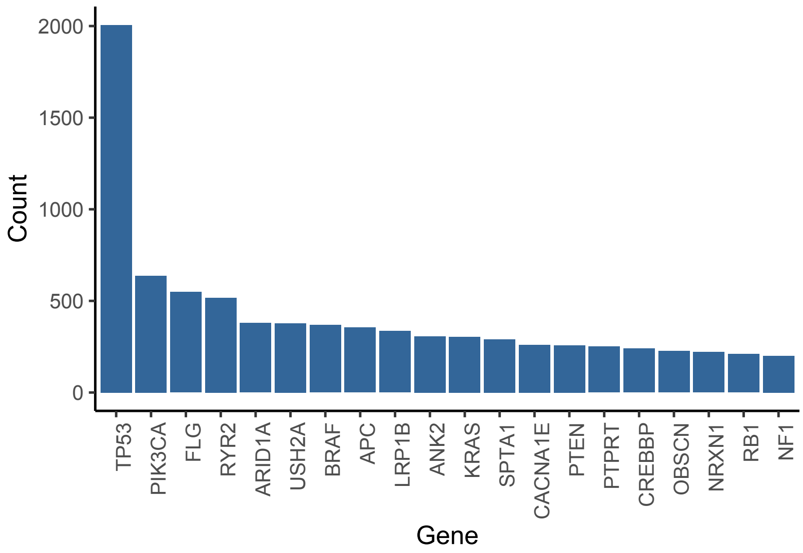

**Supplementary Fig. 7.** Density plots showing distribution of β values across all cacer types in each regulatory region.

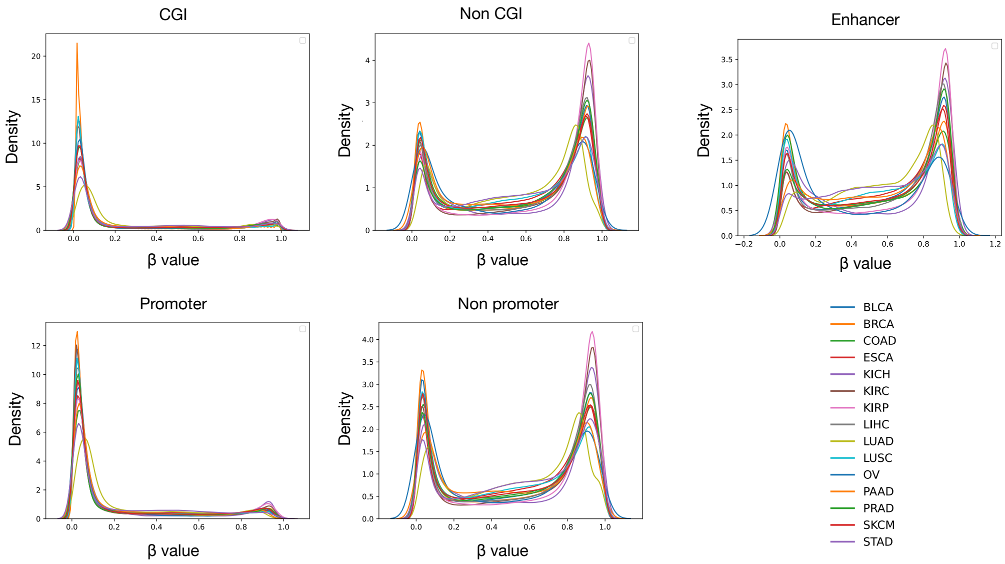

**Supplementary Fig. 8**. t-SNE plots showing the low-dimensional output of decoded methylation values from the autoencoders across all cancer types in varying regulatory regions. Color label indicates ‘High’ or ‘Low’ methylation status based on the mean methylation β values along each region.

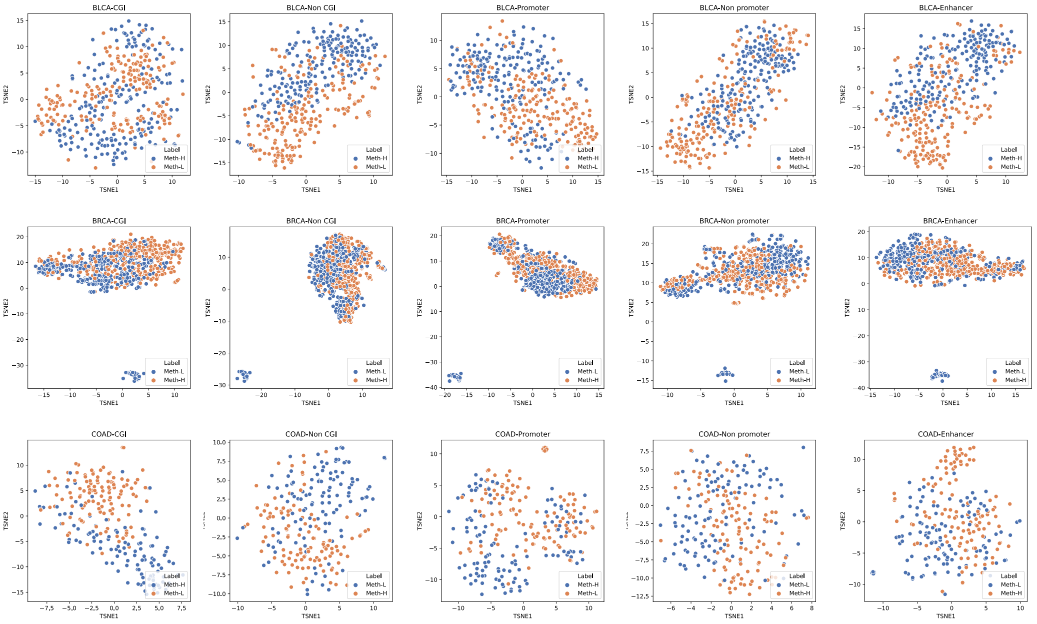

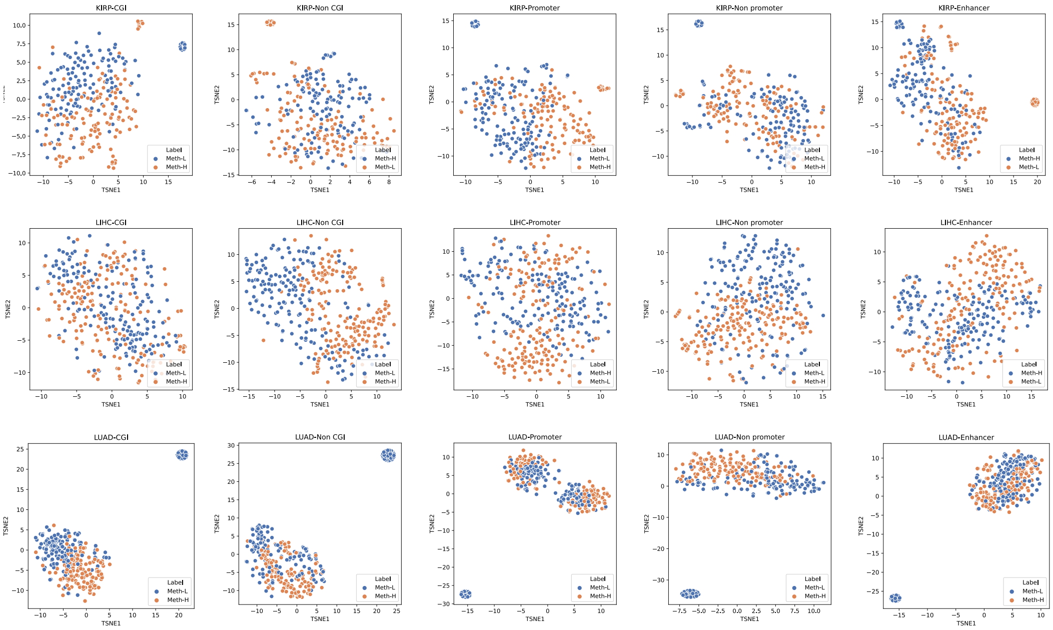
KIRP-CGI
KIRP-Non CGI
KIRP-Promoter
KIRP-Non promoter
KIRP-Enhancer
LIHC-CGI
LIHC-Non CGI
LIHC-Promoter
LIHC-Non promoter
LIHC-Enhancer
LUAD-CGI
LUAD-Non CGI
LUAD-Promoter
LUAD-Non promoter
LUAD-Enhancer
TSNE1
TSNE2
Label
Meth-L
Meth-H

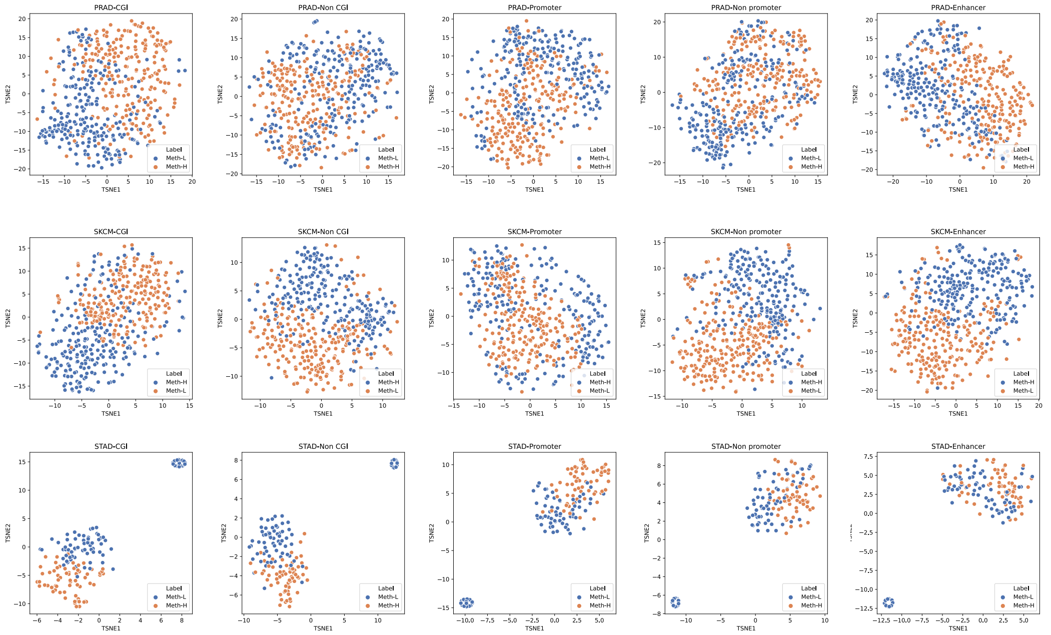
PRAD-CGI
PRAD-Non CGI
PRAD-Promoter
PRAD-Non promoter
PRAD-Enhancer
SKCM-CGI
SKCM-Non CGI
SKCM-Promoter
SKCM-Non promoter
SKCM-Enhancer
STAD-CGI
STAD-Non CGI
STAD-Promoter
STAD-Non promoter
STAD-Enhancer
TSNE1
TSNE2
Label
Meth-L
Meth-H

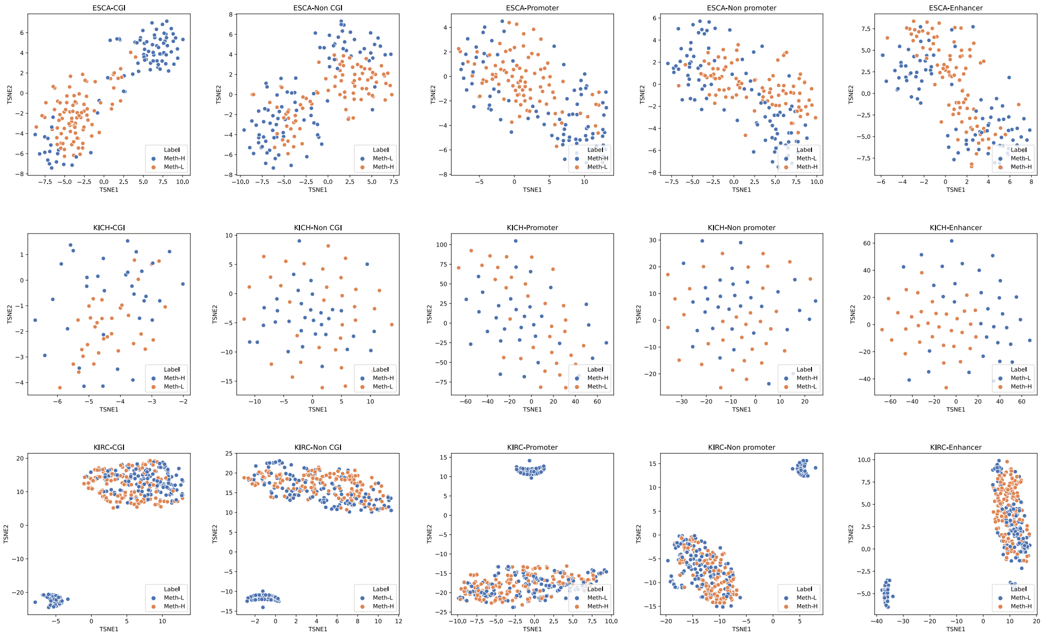

ESCA-CGI
ESCA-Non CGI
ESCA-Promoter
ESCA-Non promoter
ESCA-Enhancer
KICH-CGI
KICH-Non CGI
KICH-Promoter
KICH-Non promoter
KICH-Enhancer
KIRC-CGI
KIRC-Non CGI
KIRC-Promoter
KIRC-Non promoter
KIRC-Enhancer
TSNE1
TSNE2
Label
Meth-H
Meth-L

**Supplementary Fig. 9.** The number of undirected edges derived from STRING PPI.

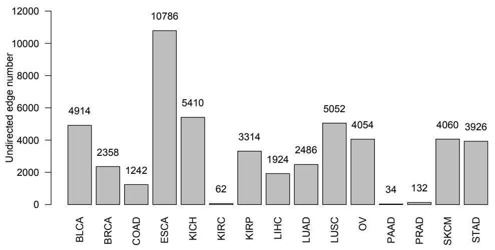


**Supplementary Fig. 10.** Thresholds of correlation coefficients used to construct final co-expression networks.

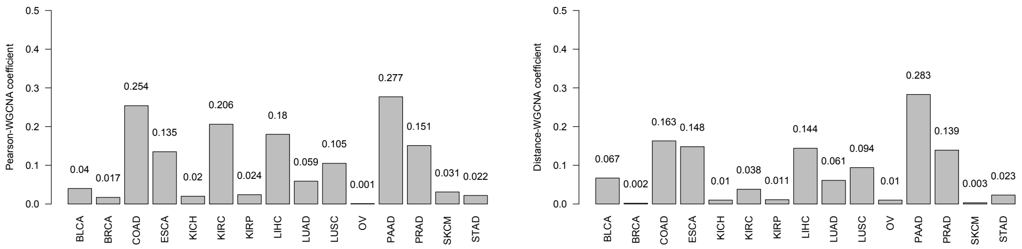

**Supplementary Fig. 11.** Training and testing accuracies for the combined models.

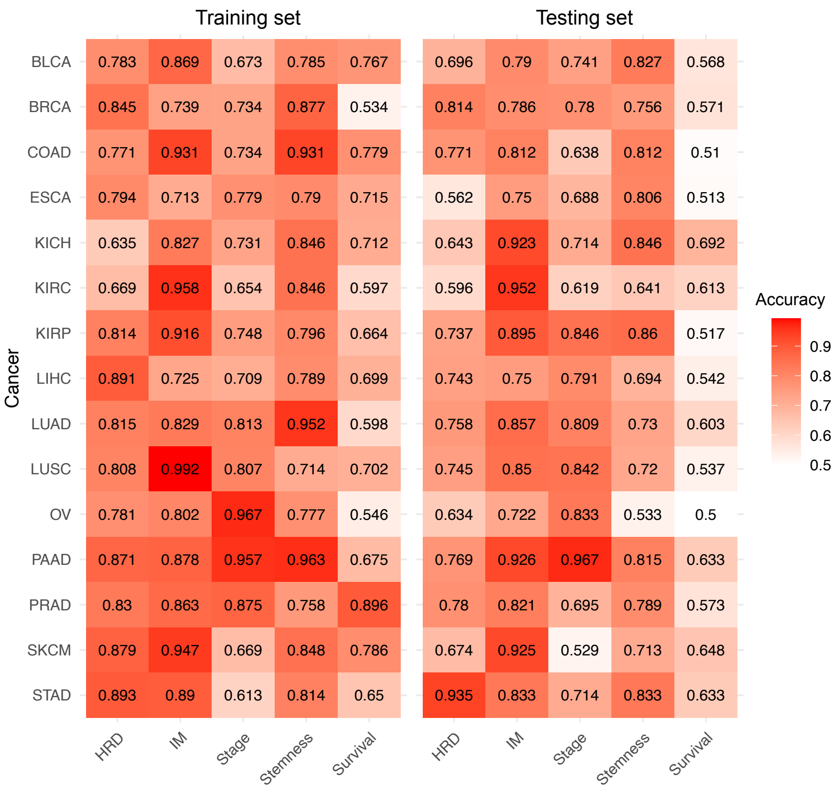

**Supplementary Fig. 12.** Testing accuracy for the top 30 combined models.

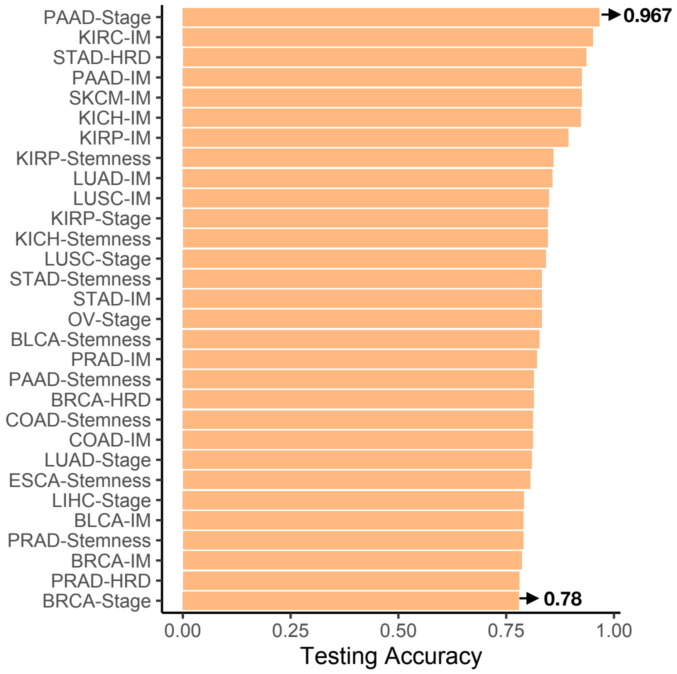

**Supplementary Fig. 13.** Cumulative node feature importance and subgraphs with overlapping CDgs derived from STRING and Distance WGCNA models. The feature index: 0=gene expression; 1=CNV; 2=mutation; 3=CpG island; 4=non-CpG island; 5=promoter; 6=non-promoter; 7=enhancer. Color bar for the subgraph indicates the normalized node importance value.

**STAD-Stemness**

**a**

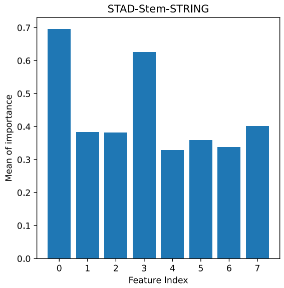


**b**

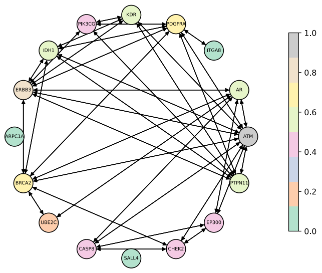


**a**

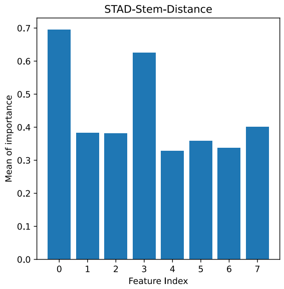


**b**

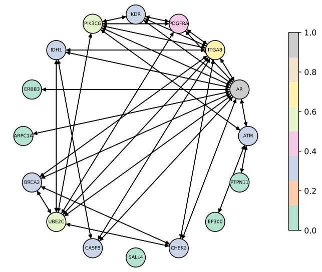

### STAD-HRD

a b

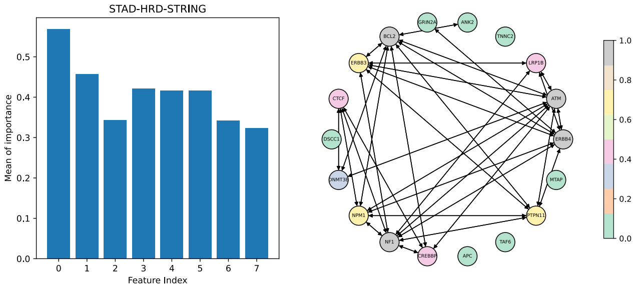


a b

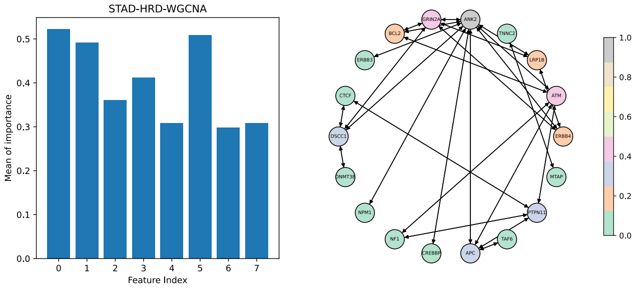


### STAD-IM

a b

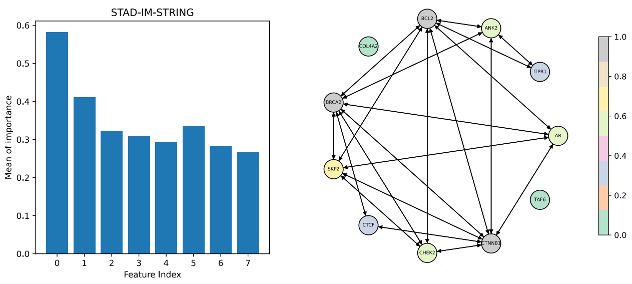


a b

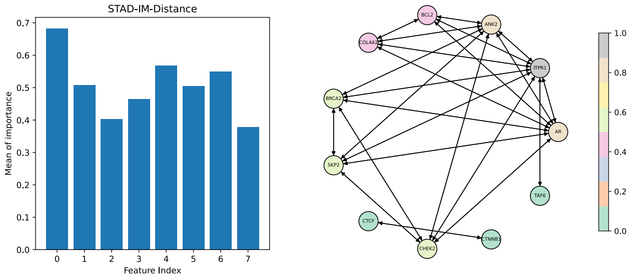

### BRCA-HRD

**a**

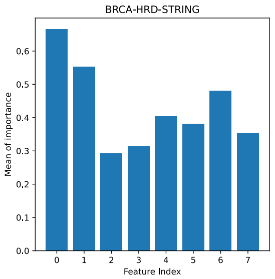


**b**

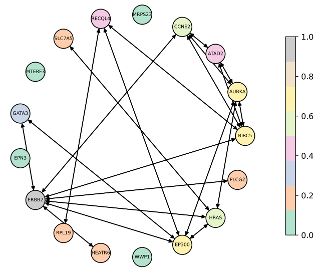

**a**

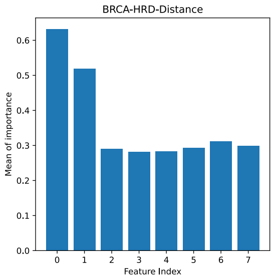


**b**

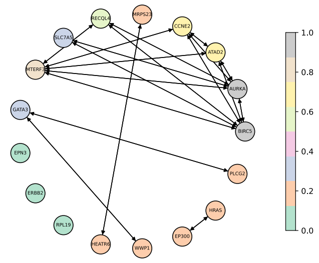

### BRCA-IM

**a**

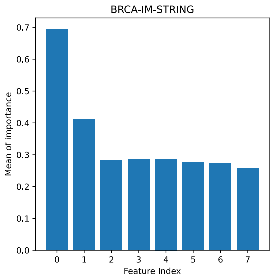


**b**

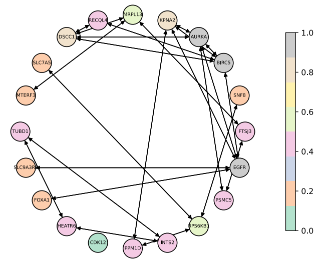

**a**

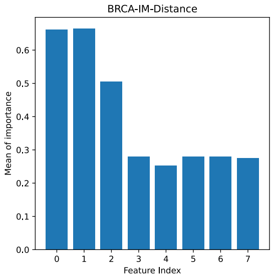


**b**

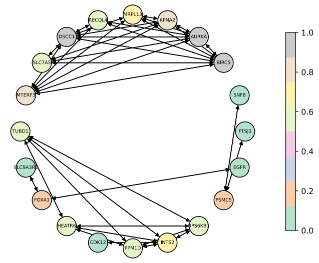

### PRAD-HRD

a

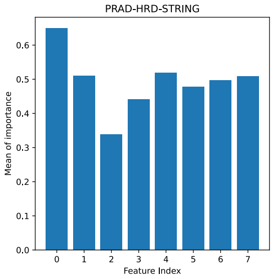


b

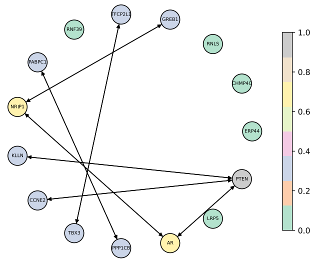


a

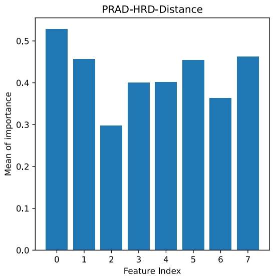


b

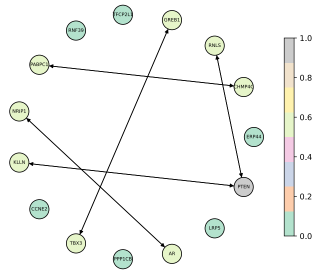


### PRAD-Stem

a

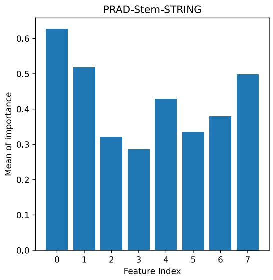


b

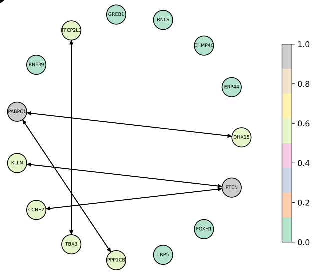


a

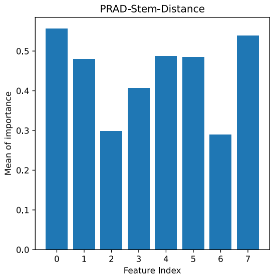


b

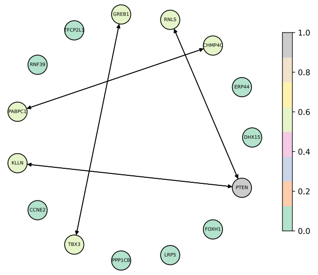

### PAAD-IM

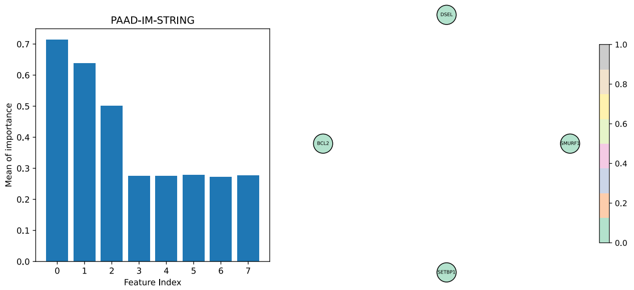


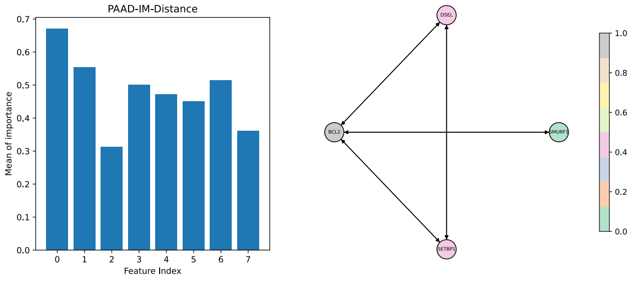


### PAAD-Stem

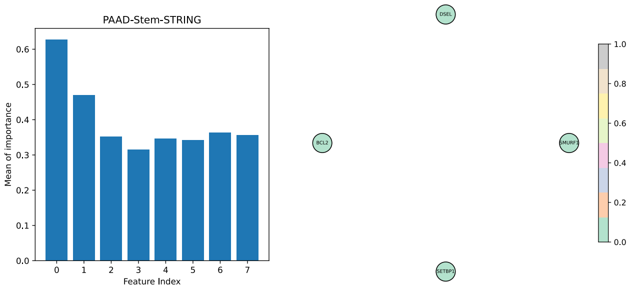


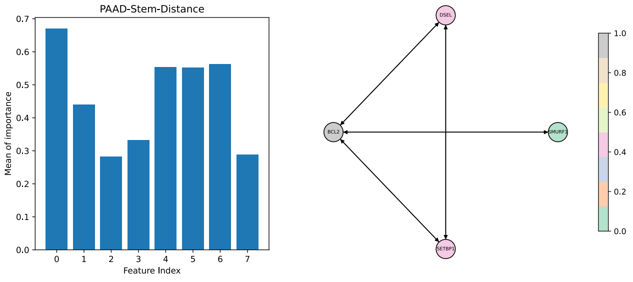

### KIRP-Stem

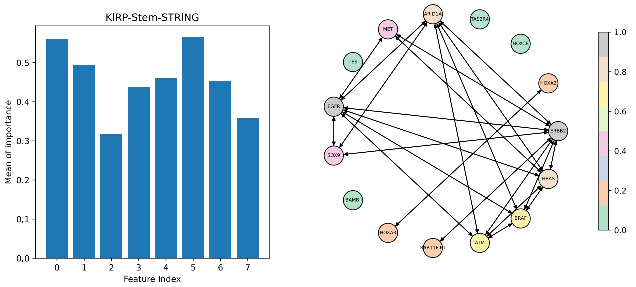


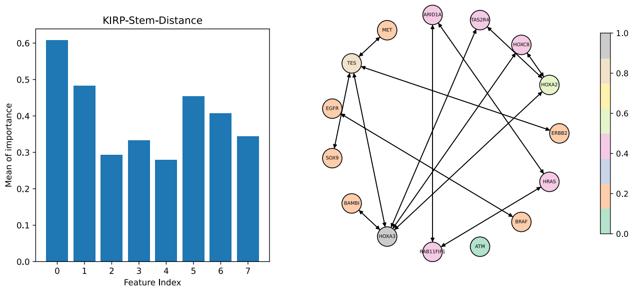


### BLCA-Stem

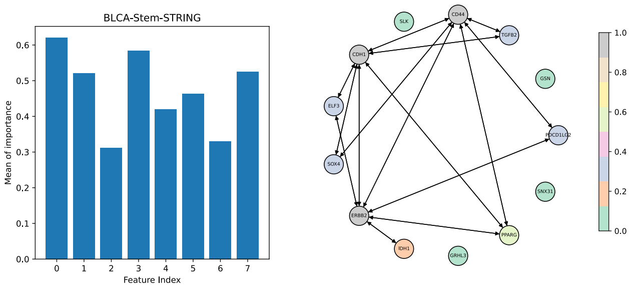


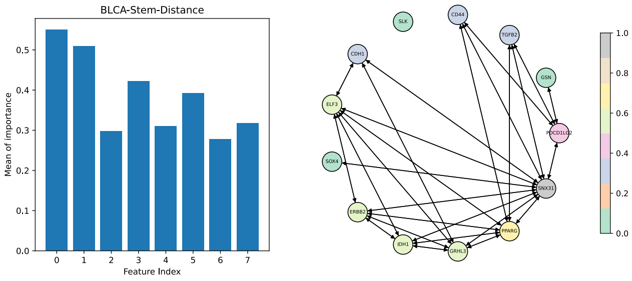

### BLCA-IM

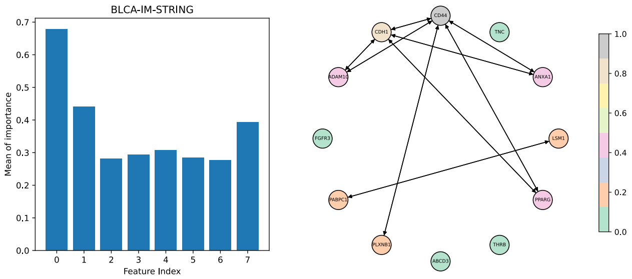


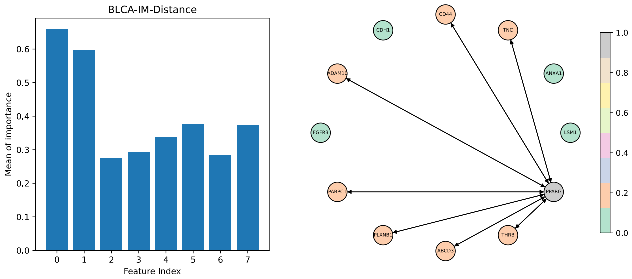


### ESCA-Stem

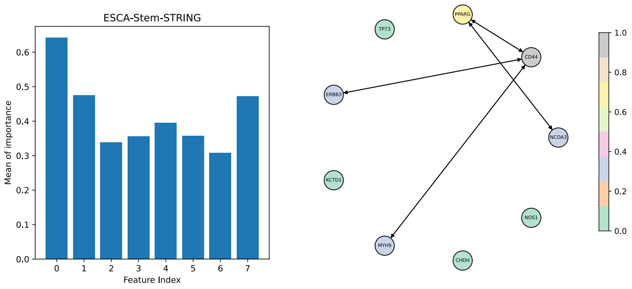


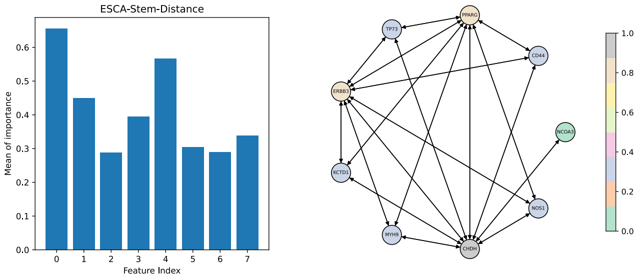

#### COAD-Stem

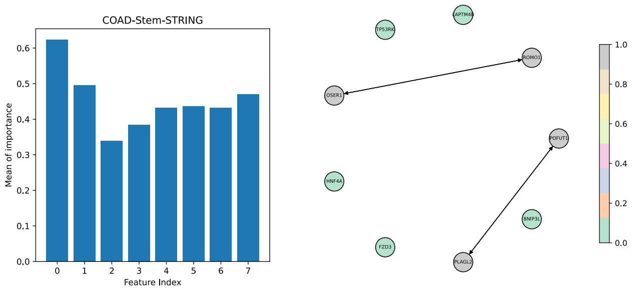


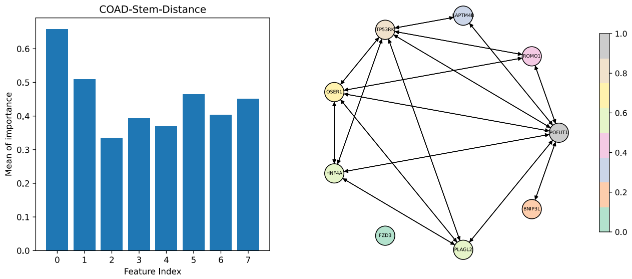


#### COAD-IM

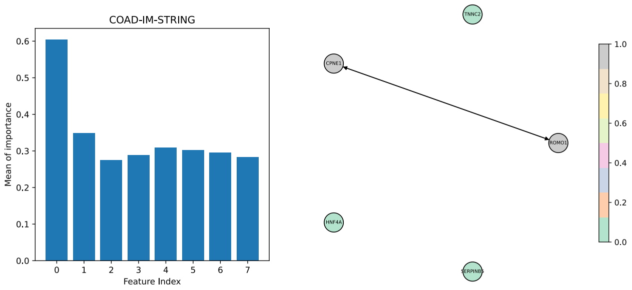


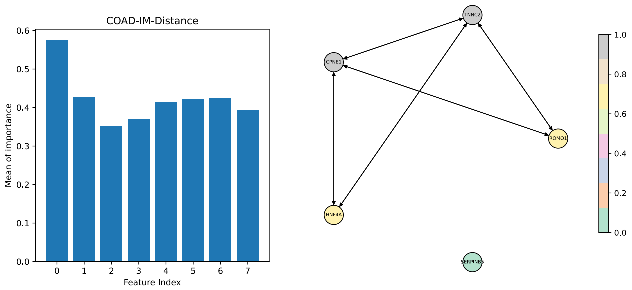

### SKCM-IM

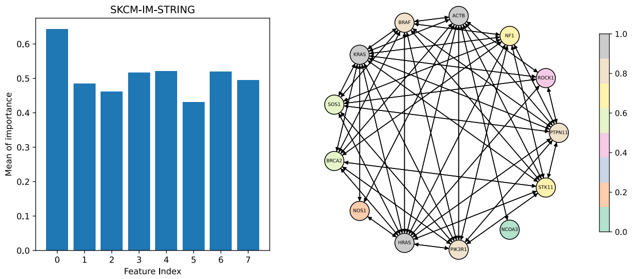


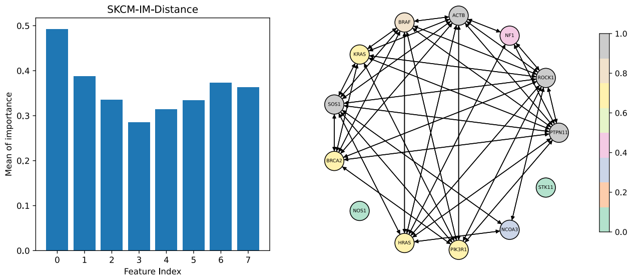


### KICH-IM

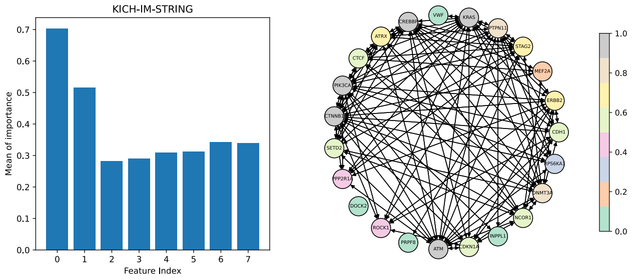


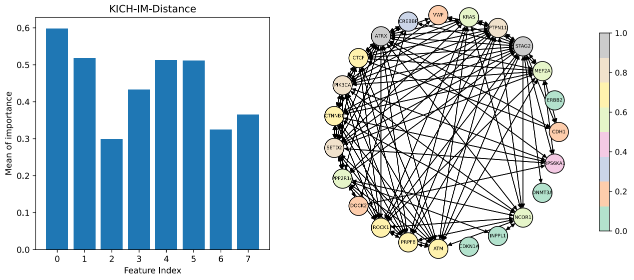
